# TNFAIP8 is a novel phosphoinositide-binding inhibitory regulator of Rho GTPases that promotes cancer cell migration

**DOI:** 10.1101/2021.03.26.437243

**Authors:** Mei Lin, Honghong Sun, Svetlana A. Fayngerts, Peiwei Huangyang, Youhai H. Chen

## Abstract

More than half of human tumors exhibit aberrantly dysregulated phosphoinositide signaling, yet how this is controlled remains not fully understood. While somatic mutations of PI3K, PTEN and Ras account for many cases of the hyperactivated lipid signals, other mechanisms for these dysfunctions in cancer are also being discovered. We report here that TNFAIP8 interacts with PtdIns(4,5)*P*_2_ and PtdIns(3,4,5)*P*_3_ and is likely to be hijacked by cancer cells to facilitate directional migration during malignant transformation. TNFAIP8 maintains the quiescent cellular state by sequestering inactive Rho GTPases in the cytosolic pool, which can be set free upon chemoattractant activation at the leading edge. Consequently, loss of TNFAIP8 results in severe defects of chemotaxis and adhesion. Thus, TNFAIP8, whose expression can be induced by inflammatory cytokines such as TNF*α* from tumor microenvironment, represents a molecular bridge from inflammation to cancer by linking NF-κB pathway to phosphoinositide signaling. Our study on the conserved hydrophobic cavity structure will also advise *in silico* drug screening and development of new TNFAIP8-based strategies to combat malignant human diseases.

## INTRODUCTION

Cancer cell locomotion is an important form of eukaryotic cell migration. Failure to treat cancer metastases from solid tumors in clinic has been responsible for the majority of patient deaths^1^. The cascades for metastasis are complicated and encompass migration of cancerous cells away from the primary site, intravasation into the circulation system, extravasation to secondary tissues, and formation of distant metastatic tumors^2^. Cell migration is a pivotal step during these metastatic processes. In order to move in one direction, the cells must first form a defined front and rear. This polarity is characterized by asymmetrical activation of proteins such as PI3Ks, Rho GTPases, and actin regulatory proteins at the leading and trailing edges. Despite intensive studies, how the shallow gradients of chemoattractants translate into axes of cell polarization during chemotaxis is not well understood. Phosphatidylinositol (PtdIns) is a unique membrane phospholipid that can be phosphorylated at the 3, 4 and 5 positions of the inositol ring to generate seven phosphoinositides, among which PtdIns(4,5)*P*_2_ (PIP2) and PtdIns(3,4,5)*P*_3_ (PIP3) are important lipid second messengers^3^. PtdIns(4,5)*P*_2_ is the most abundant phosphoinositide and binds to proteins important for actin polymerization, focal contacts formation, and cell adhesion. PtdIns(3,4,5)*P*_3_ is highly enriched in the leading edge and controls cell migration by promoting actin polymerization through both GTPase-dependent and independent mechanisms. In addition, PI3K-mediated generation of PtdIns(3,4,5)*P*_3_ and the subsequent activation of Akt and its downstream cascades (e.g., mTORC1) compose the signaling axis controlling cancer cell survival and growth. Rho small GTPases are conformationally regulated by the binding of GTP and GDP, and they are activated by guanine nucleotide exchange factors (GEFs) and inactivated by GTPase activating proteins (GAPs). Moreover, Rho family can bind to guanine nucleotide dissociation inhibitors (GDIs), which consist of RhoGDIα (ubiquitously expressed), RhoGDIβ (hematopoietic tissue-specific expression), and RhoGDIγ (preferentially expressed in brain, kidney, lung, pancreas, and testis)^4^. Of the at least 20 Rho family proteins identified so far in human, Rac1/2, Cdc42, and RhoA/B have been the most widely studied for their importance in cell migration. Rac and Cdc42 are mainly active at the leading edge, contributing to spatially confined actin polymerization and protrusion via Arp2/3 and WASP/WAVE dependent mechanisms, while Rho is involved in rear retraction^5^. It remains to be characterized how these molecular machineries are coordinated temporally and spatially to enable cells to sense chemokine gradients, induce polarized morphology, and rapidly reorganize the cytoskeleton.

The TIPE (tumor necrosis factor-*α*-induced protein 8 (TNFAIP8)-like, or TNFAIP8L) family of proteins have been implicated in inflammation and tumorigenesis. There are four highly homologous mammalian TIPE family members: TNFAIP8, TIPE1 (TNFAIP8L1), TIPE2 (TNFAIP8L2), and TIPE3 (TNFAIP8L3). Existing evidences support that TNFAIP8 is an oncogene, which is found to be frequently over-expressed in a wide range of human cancers and promote tumor metastasis^6–14^. TIPE2 is almost exclusively expressed in hematopoietic cells and is initially identified as one of the most highly upregulated genes in the spinal cord of experimental autoimmune encephalomyelitis (EAE) mice^15^. TIPE2 is reported to be a negative regulator of innate and adaptive immunity and essential for maintaining immune homeostasis^16, 17^. TIPE2 can direct myeloid cell polarization^18^ and play a redundant role with TNFAIP8 in controlling murine lymphocyte migration during neural inflammation^19^. TIPE3 is preferentially expressed in secretory epithelial tissues^20^, and its expression is also markedly upregulated in cervical, colon, lung, and esophageal cancers^21^. The high-resolution crystal structures of murine TNFAIP8 C165S mutant, human TIPE2, and human TIPE3 have been recently resolved^21–23^, and they reveal a conserved TIPE homology (TH) domain composed of a large hydrophobic central cavity. Although the outer surface is highly charged, the central cavity is predominantly hydrophobic, implicating lipid binding capacity. In this report, we describe TNFAIP8 as a phosphoinositide-binding negative regulator of Rho GTPases and examine its role in cancer cell migration.

## RESULTS

### Defective chemotaxis and adhesion in TNFAIP8-deficient differentiated HL-60 cells

TIPE proteins are newly discovered risk factors for inflammation and cancer (**Supplementary Fig. 1a**). TNFAIP8, the founding member of this family, is originally identified as one of the most differentially expressed genes in primary and matched metastatic head and neck squamous cell carcinoma (HNSCC) from the same patient^24, 25^. To investigate the role of TNFAIP8 in directed cell migration, we used both loss- and gain-of-function approaches to manipulate TNFAIP8 expression in HL-60 (**Fig. 1a**), a human promyelocytic leukemia cell line that can be conveniently differentiated into neutrophil-like cells *in vitro* as a powerful model for eukaryotic chemotaxis^26^. Human TNFAIP8 has six transcript variants (Variant 1 to 6) with different transcription start sites/exons and can be translated to four protein isoforms (Isoform a to d). Among these, Variants 3 to 6 were minimally expressed in normal or cancer tissues according to recent RNA-seq analysis^27^, whereas Variant 2 was frequently over-expressed and Variant 1 was commonly downregulated in multiple human malignancies. By targeting the shared exon with CRISPR/Cas9, we validated the complete knockout of all TNFAIP8 isoforms in the established HL-60 clonal cell lines (**Supplementary Fig. 1b–d**). To investigate the effects of TNFAIP8 deficiency in differentiated HL-60 (dHL-60) cells, we first tested chemoattractant EC50 and verified that 10 nM fMLP was optimal for migration with an around eight-fold increase in cell numbers through the Transwell filters (**Supplementary Fig. 2a**), while CXCL8 induced a two-fold increase within a wide range of concentrations (**Supplementary Fig. 2b**). To demonstrate whether TNFAIP8 was necessary and sufficient in regulating dHL-60 chemotaxis, TNFAIP8 expression was rescued by lentiviral titration and infection into knockout cells to the endogenous level (**Fig. 1a**). We found that TNFAIP8 knockout (TKO) dHL-60 cells displayed substantial transmigration defects towards fMLP and CXCL8 but did not have obvious defects in random migration (without chemoattractants), relative to those of wild-type (WT) counterparts. These behaviors were consistent on either bare Transwell membranes (**Fig. 1b**) or Matrigel coated filters (**Fig. 1c**). Furthermore, the impaired chemotaxis could be completely ‘rescued’ by re-expressing a wild-type TNFAIP8 transgene (**Fig. 1b**,**c**). These results indicate that TNFAIP8 is essential for the chemotaxis of human dHL-60 cells *in vitro*.

**Figure 1.**
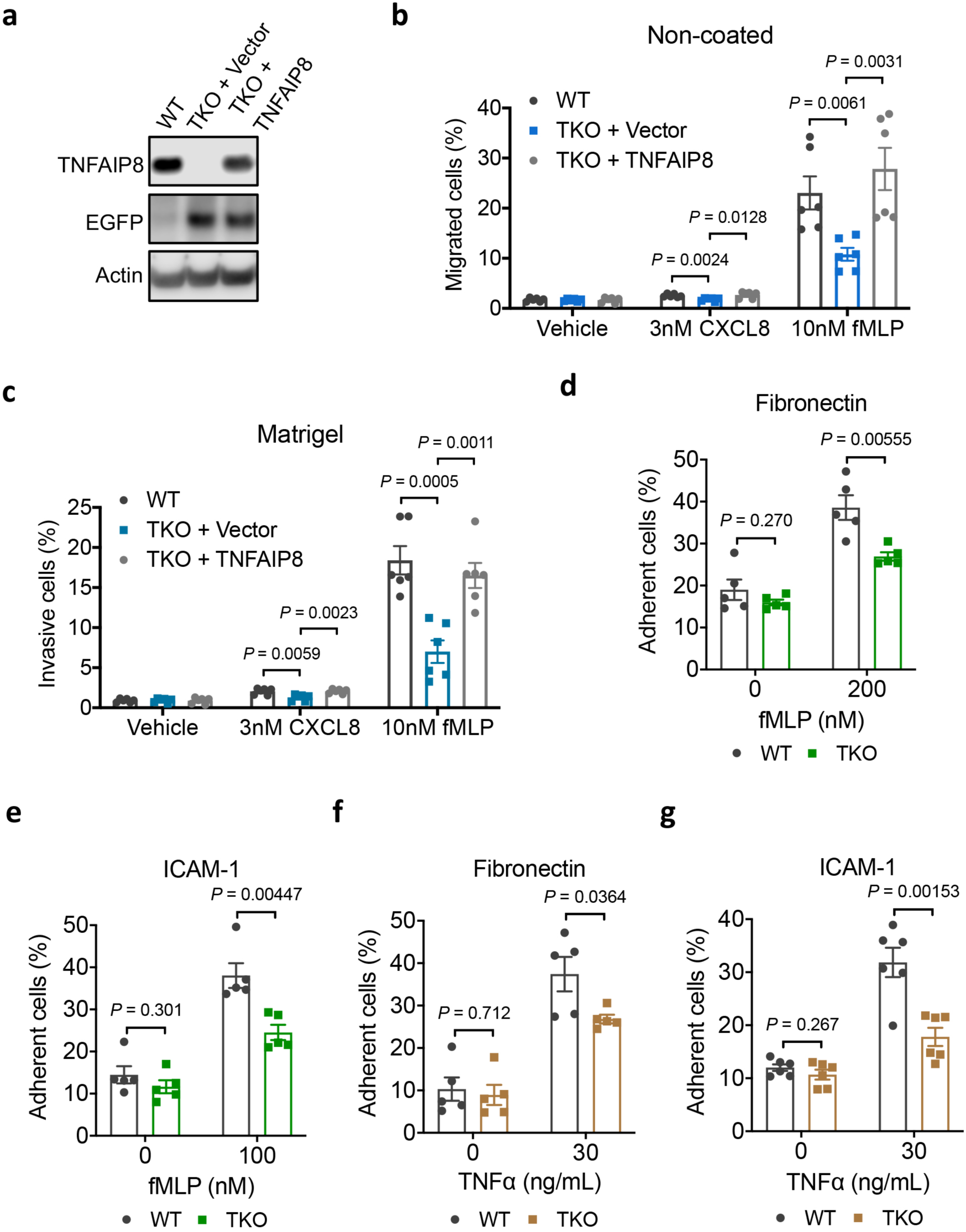
TNFAIP8 is required for chemotaxis and adhesion of neutrophil-like differentiated HL-60 (dHL-60) cells *in vitro*. (**a**) Protein levels of TNFAIP8 and EGFP in wild-type (WT), TNFAIP8 knockout (TKO), and lentivirally rescued TNFAIP8-expressing dHL-60 cells determined by western blot. (**b**,**c**) Migration of WT and TKO dHL-60 cells through non-coated (**b**) or Matrigel coated (**c**) Transwell filters towards 10 nM fMLP or 3 nM CXCL8 (*n* = 6 individual WT or TKO clonal cell lines established from CRISPR/Cas9). (**d**–**g**) WT and TKO dHL-60 cells were serum starved for 2 h and subsequently stimulated with 100–200 nM fMLP (**d**,**e**) or 30 ng/mL TNFα (**f**,**g**). Percentages of adherent cells on fibronectin (**d**,**f**) or ICAM-1 (**e**,**g**) coated surfaces were calculated relative to unwashed wells representing 100% (*n* = 5 (**d**–**f**) or 6 (**g**) clones). Data represent three (**a**–**g**) independent experiments with similar results (mean ± s.e.m.). *P* values are from two-sided unpaired Student’s *t*-test (**b**–**g**).

Cell adhesion plays a critical role in the extravasation of leukocytes and tumor cells in inflamed tissues, and the ability of cells to regulate adhesion derives from their ability to control the affinity states of integrins. Integrin ‘inside-out’ signaling is the process where signals inside the cells cause the integrin external domains to assume an activated state^28^. Integrin affinity can be activated by several signaling events, including G protein coupled receptor (GPCR) activation as well as chemokine and cytokine stimulation, which allow cells to undergo arrest resulting in firm adhesion on endothelia expressing intercellular adhesion molecules (ICAMs). To test whether TNFAIP8 deficiency also affected adhesive capacity, WT and TKO cells were serum starved and subsequently stimulated with fMLP or TNFα before being placed on fibronectin or ICAM-1 coated surfaces. Notably, TKO dHL-60s exhibited significantly reduced adhesion to fibronectin (**Fig. 1d,f**) or ICAM-1 (**Fig. 1e,g**) compared to their WT counterparts when treated with either fMLP (**Fig. 1d,e**) or TNFα (**Fig. 1f,g**). In addition, they didn’t significantly differ when no chemokine or cytokine was present. These results indicate that TNFAIP8 knockout confers deficiency in ECM adhesiveness.

### Reduced growth and survival of TNFAIP8-deficient HL-60 cells

We observed that TKO HL-60s had reduced proliferation in culture media containing either 1% FBS (**Fig. 2a**) or 10% FBS (**Fig. 2b**) compared to WT cells. Additionally, cell counting with trypan blue exclusion revealed a lower viability in TKO HL-60s after two-day culture in 1% FBS media relative to WT cells (**Fig. 2c**). These results suggest that TNFAIP8 deficiency can decrease HL-60 cell growth and survival *in vitro*. Moreover, TNFAIP8 knockout in HL-60 cells considerably attenuated the phosphorylation of ERK1/2 (T202/Y204) and p70S6K (T389 and S371), both of which can regulate cyclin D1 levels and consequently G1-S transition. In contrast, the phosphorylation of LIMK (LIMK1 T508/LIMK2 T505), cofilin (S3), p38 (T180/Y182), 4E-BP1 (T37/46) were not influenced by TNFAIP8 in undifferentiated cells (**Fig. 2d**). These results suggest that TNFAIP8 may play a role in the regulation of PI3K-AKT-mTOR and MEK-ERK signaling pathways in proliferating HL-60s.

**Figure 2.**
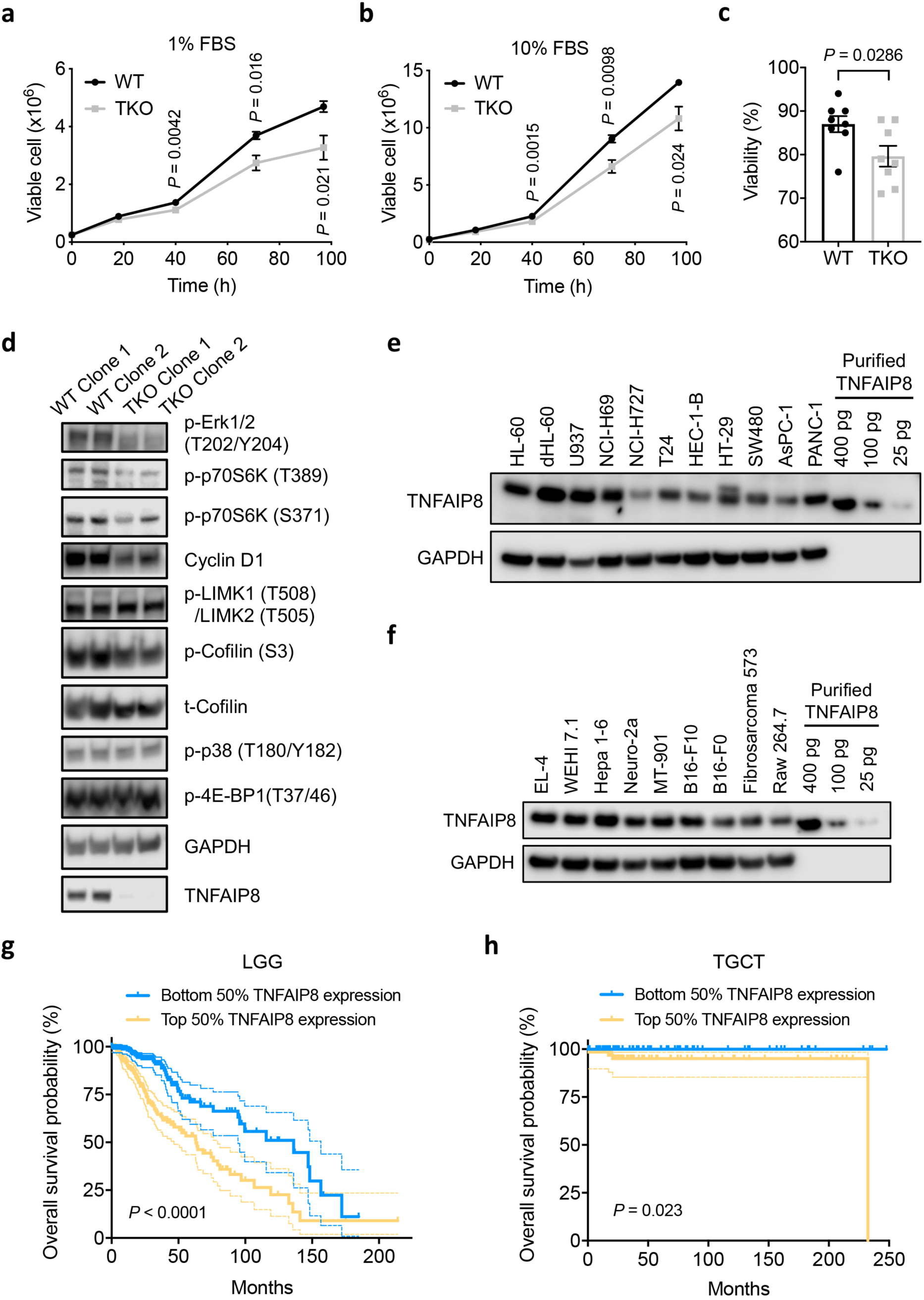
TNFAIP8 deficiency decreases the proliferation and survival of HL-60 cells *in vitro*, and high TNFAIP8 expression is associated with poor prognosis in human cancers. (**a**,**b**) Viable cell numbers of WT and TKO HL-60 in culture media containing 1% FBS (**a**) or 10% FBS (**b**) measured by CellTiter-Glo (CTG) assay over the indicated time (*n* = 4 clones). (**c**) Cellular viability after two-day culture in 1% FBS containing media detected by trypan blue exclusion assay (*n* = 8 clones). (**d**) WT and TKO HL-60 cells were analyzed by western blot using antibodies against the indicated proteins. (**e**,**f**) Immunoblot analysis of TNFAIP8 protein levels in various human (**e**) and mouse (**f**) cancer cell lines. GAPDH served as a loading control. Purified human TNFAIP8 isoform b protein was used for quantification. (**g**,**h**) Kaplan–Meier survival curves of 510 brain lower grade glioma (LGG) patients (**g**) or 134 testicular germ cell tumor (TGCT) patients (**h**) enrolled in TCGA database. Patients were equally divided into two groups (top and bottom 50% TNFAIP8 expression) based on TNFAIP8 mRNA levels in their tumors. Dashed lines indicate the 95% confidence intervals. Data represent three (**a**–**c**) or two (**d**–**f**) independent experiments with similar results (mean ± s.e.m.). *P* values are from two-sided unpaired Student’s *t*-test (**a**–**c**) or log-rank (Mantel–Cox) test (**g**,**h**).

### Association between TNFAIP8 expression and poor prognosis in human cancers

High TNFAIP8 expression has been detected in various clinical specimens compared to healthy tissues. Studies have reported TNFAIP8 as a risk factor for the progression and poor prognosis of numerous human malignancies, including prostate cancer^8^, gastric adenocarcinoma^6, 9, 10, 29^, esophageal squamous cell carcinoma (ESCC)^13^, non-small-cell lung cancer^7^, and non-Hodgkin’s lymphoma^11^. We quantified TNFAIP8 protein expression in numerous human (**Fig. 2e**) and mouse (**Fig. 2f**) cancer cell lines and found that TNFAIP8 protein level had a 2.5-fold increase after HL-60 differentiation, which made dHL-60 the most highly expressing cell line among all cancer types examined. In addition, higher TNFAIP8 mRNA expression correlated significantly with worse prognosis in patients with brain lower grade glioma (LGG) or testicular germ cell tumor (TGCT) based on TCGA analysis (**Fig. 2g,h**). To gain insights on subcellular localization of human TNFAIP8, we fractionated proteomic samples from several cancer cell lines into cytosol, membrane, nuclear, and cytoskeleton components (**Supplementary Fig. 2c–e**). Notably, TNFAIP8 protein localized exclusively in the cytoplasm under resting conditions in HL-60 leukemia cells (10.2 ± 2.0 pg TNFAIP8 per μg total protein), U-937 lymphoma cells (15.7 ± 0.4 pg TNFAIP8 per μg total protein), and NCI-H69 small cell lung carcinoma cells (12.3 ± 1.0 pg TNFAIP8 per μg total protein).

### Aberrant phosphoinositide levels resulted from TNFAIP8 deficiency in quiescent cells

During chemotaxis cells are polarized, with a filamentous actin-rich lamellipodium at the front and uropod at the rear^30^. Localized accumulation of phosphatidylinositol lipids has been proposed to be the key event directing the recruitment and activation of signaling components required for chemotaxis. Upon chemoattractant stimulation, PI3K is locally activated by ligand-bound GPCRs or receptor tyrosine kinases (RTKs). Active PI3K catalyzes phosphorylation at the 3 position of PtdIns(4,5)*P*_2_ to generate PtdIns(3,4,5)*P*_3_, which leads to its enrichment at the leading edge. The asymmetric distribution of PtdIns(3,4,5)*P*_3_ is also confined spatially by the actions of 3’ lipid phosphatases, which convert PtdIns(3,4,5)*P*_3_ back to PtdIns(4,5)*P*_2_; phosphatase and tensin homolog on chromosome ten (PTEN) is enriched at the back and sides of migratory cells. To explore if there were changes in total phosphoinositide levels in quiescent and stimulated dHL-60 cells, we performed protein-lipid overlay assay with GST-GRP1-PH (PtdIns(3,4,5)*P*_3_ sensor), GST-PLCδ-PH (PtdIns(4,5)*P*_2_ sensor), and GST-SidC-3C (PtdIns(4)*P* sensor) proteins. Notably, we detected markedly increased total PtdIns(3,4,5)*P*_3_ and PtdIns(4)*P*, but reduced PtdIns(4,5)*P*_2_ level in TNFAIP8-deficient dHL-60s under resting condition (**Fig. 3a–d**). By contrast, the differences diminished after 60 s of fMLP treatment and the cellular levels of these phosphoinositides became comparable in WT and TKO cells (**Fig. 3a–d**). Additionally, TKO dHL-60 cells displayed increased cell number and survival compared to WT cells after inducing differentiation for 5 days (**Supplementary Fig. 2f,g**), which was likely due to the upregulated PtdIns(3,4,5)*P*_3_ level in these cells. Consistently, we also observed significant increases of total PtdIns(3,4,5)*P*_3_ in TNFAIP8 and TIPE2-deficient (DKO) murine splenocytes (**Supplementary Fig. 3a**) and thymocytes (**Supplementary Fig. 3b**), where TIPEs were known to have extremely high expressions. These results suggest that TNFAIP8 can regulate phosphoinositide metabolism and signaling.

**Figure 3.**
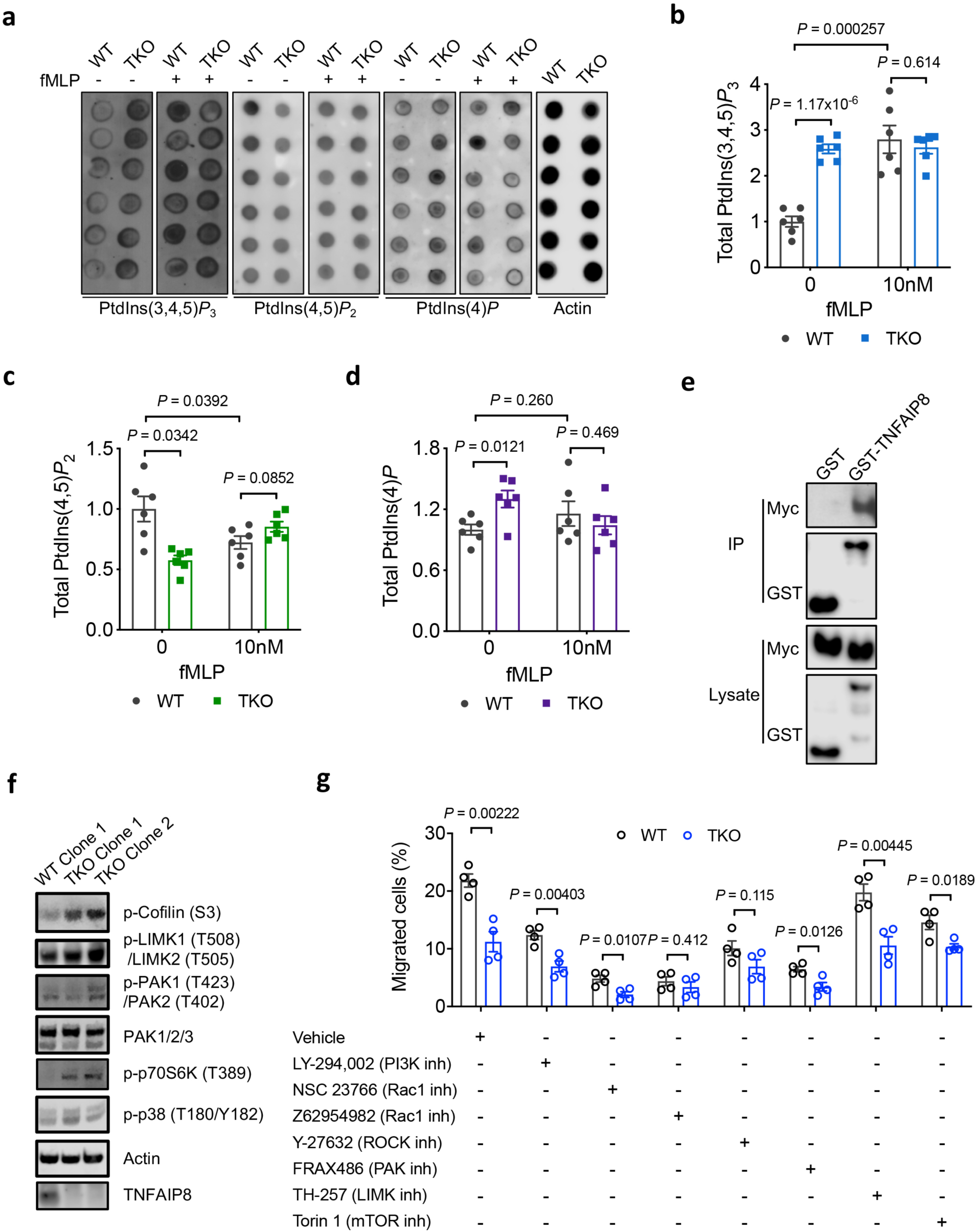
TNFAIP8 regulates cellular phosphoinositide levels and Rho GTPase signaling pathways. (**a**–**d**) Protein-lipid overlay assay (**a**) showing cellular levels of PtdIns(3,4,5)*P*_3_ (**b**), PtdIns(4,5)*P*_2_ (**c**), and PtdIns(4)*P* (**d**) in WT and TKO dHL-60s examined with GST-GRP1-PH, GST-PLCδ-PH, or GST-SidC-3C protein, respectively. dHL-60 cells were stimulated with DMSO vehicle or 10 nM fMLP at 37°C for 60 s. Signal from unstimulated WT dHL-60 cells was set as 1 (*n* = 6 clones). (**e**) Immunoblot analysis of the indicated proteins in 293T cells expressing Myc-tagged Cdc42 along with GST or recombinant GST-tagged TNFAIP8 before (lower panel) or after (upper panel) pull-down with glutathione beads. (**f**) WT and TKO dHL-60 cells were analyzed by western blot using antibodies against the indicated proteins. (**g**) Migration of WT and TKO dHL-60s across 3 μm pore size non-coated Transwell filters towards 10 nM fMLP, with or without pretreatment with the indicated inhibitors (*n* = 4 clones). Data represent at least three (**a**–**d**) or two (**e**–**g**) independent experiments with similar results (mean ± s.e.m.). *P* values are from two-sided unpaired Student’s *t*-test (**b**–**d**,**g**).

### Regulation of Rho GTPase-dependent signaling by TNFAIP8

The directed migration of cells requires membrane protrusion, attachment to the substratum, and release of the cell rear followed by retraction of the tail. Phosphoinositides including PtdIns(4,5)*P*_2_ and PtdIns(3,4,5)*P*_3_ are major second messengers relaying chemotaxic signals from receptors and exert their functions through interacting with PIP effectors such as pleckstrin homology (PH) domain and polybasic motif (PBM) proteins. A subset of PH-proteins includes GEFs and GAPs for the Rho family. During migration, active Rac-GTP preferentially localizes at the front of the cell, while at the trailing uropod Rac is inactive, which prevents the formation of multiple lamellipodia^4^; this front-to-rear polarity is essential for directional migration. Rac-GTP promotes actin polymerization through effector proteins such as p21-activated kinases (PAKs) and WASP-family verprolin-homologous protein (WAVE). Cdc42 can bind and activate the Par polarity complex, which could in turn activate Rac through T-cell lymphoma invasion and metastasis-inducing protein 1 (Tiam1)^31^. TIPE2 was reported to co-immunoprecipitate with Rac in 293T cells or dHL-60s stimulated with fMLP^18^. Moreover, TIPE2 was able to inhibit Rac membrane translocation/activation and suppress Rac downstream signaling via direct binding to the hydrophobic C-terminal CAAX motif region of Rac^16^. To explore the interactions between TNFAIP8 and Rho family of small GTPases, we assessed immunoblot of TNFAIP8 and Cdc42 in 293T cells expressing recombinant GST-tagged TNFAIP8 and Myc-tagged Cdc42 following GST pull-down. Myc-Cdc42 was detected to immunoprecipitate with GST-TNFAIP8 (**Fig. 3e**). Consistently, we also found that TIPE2 associated with Myc-tagged Cdc42 (**Supplementary Fig. 3c**) or Par3 (**Supplementary Fig. 3d**) but not Par6 (data not shown) when GST-tagged TIPE2 was expressed and pulled down in 293T cells. In addition, immunoprecipitation with anti-TIPE2 antibody and immunoblot for AKT revealed their endogenous association in Raw 264.7 macrophages (**Supplementary Fig. 3e**). Furthermore, TIPE2 preferentially associated with the Cdc42-17N mutant (GDP-bound form) rather than the Cdc42-61L mutant (GTP-bound form) (**Supplementary Fig. 3f**). These results indicate that TIPEs may function through Rho GTPases including Rac and Cdc42, as well as PI3K-AKT pathway to contribute to chemotaxic actin remodeling.

To further explore the mechanisms by which TNFAIP8 regulates directional migration, we performed western blot and found that TNFAIP8-deficient dHL-60s exhibited upregulated phosphorylated cofilin (S3), LIMK (LIMK1 T508/LIMK2 T505), PAK (PAK1 T423/PAK2 T402) as well as p70S6K (T389) signals (**Fig. 3f**). By contrast, no difference was observed in the phosphorylation of p38 (T180/Y182) MAP kinase pathway between WT and TKO groups (**Fig. 3f**). We further manipulated the molecular activities using various pharmacological blockers (**Fig. 3g**). The transmigration of both WT and TKO cells could be attenuated markedly by the antagonists LY-294,002 (PI3K inhibitor), NSC 23766 (selective inhibitor of Rac1-GEF interaction), Torin1 (mTOR inhibitor XI), InSolution Z62954982 (Rac1 inhibitor II), Y-27632 (ROCK inhibitor), and FRAX486 (PAK inhibitor). Moreover, NSC 23766, InSolution Z62954982, Y-27632, FRAX486, and Torin1 effectively reduced the difference between WT and TKO groups (**Fig. 3g**), implicating the importance of these molecular pathways in TNFAIP8 modulation of dHL-60 cell migration.

### TNFAIP8 interaction with phosphoinositides through TH domain and *α*0 helix

TNFAIP8 consists of 7 *α* helixes forming a pocket that measures 20 Å in depth and 7–10 Å in diameter (**Fig. 4a** and **Supplementary Video 1**). We compared the protein topographies of TIPE family and RhoGDI1 from Protein Data Bank (PDB) by CASTp 3.0, which analytically identified and gauged the surface accessible pockets as well as interior inaccessible cavities^32^. The area and volume of TNFAIP8 C165S cavity was 579.8 and 406.0 SA (solvent accessible surface), respectively (**Fig. 4b**), exhibiting considerably larger space than that of RhoGDI1, which was known to accommodate lipid modifications of cargo proteins. According to a previous protein-lipid overlay assay, TIPEs predominantly interacted with PtdIns(4,5)*P*_2_, PtdIns(3,5)*P*_2_, PtdIns(3,4)*P*_2_, and PtdIns(3,4,5)*P*_3_ out of the 15 eukaryotic lipids screened^21^. To gain more insights into the binding modes for various phospholipids, we performed molecular docking using Glide XP docking method (**Supplementary Table 3**) and Prime MM-GBSA scoring function (**Supplementary Table 4**), which had consistency with the structural proposition that the fatty acid chains of phosphoinositides were positioned inside the TH domain pocket formed by *α*1–*α*6 helixes, while the negatively charged phosphate head was accommodated by electrostatic interactions with the positively charged amino acids located at the entrance of the cavity as well as the more flexible *α*0 helix. To further explore the phosphoinositide interactions in the TH domain region, we employed molecular dynamics (MD) simulations to elucidate the binding site and time dynamics of interacting amino acid residues. After initial structural adaptations that may be due to conformational changes from the flexible *α*0 helix, the simulation system reached equilibrium, and the secondary structures stayed intact throughout the 50 ns period (**Supplementary Fig. 4**). We observed that PtdIns(4,5)*P*_2_ and PtdIns(3,4,5)*P*_3_ molecules bound in the TH domain by forming favorable hydrophobic interactions, hydrogen bonds, water bridges, and electrostatic interactions during the simulation (**Supplementary Fig. 5a–e**). Analysis of the trajectories revealed that the critical amino acids involved in maintaining stable interactions at the binding site appeared to be H86, R87, and K103, three important positively charged residues on the surface of the cavity, as well as K32 and K36 on *α*0 helix (corresponding to H76, R77, K93, K22, and K26 in human TNFAIP8 isoform b) (**Supplementary Fig. 5f,g**). A similar binding mode and interactions were also found for the phosphatidylinositol, monophosphates, and bisphosphates (**Supplementary Fig. 6**). Additionally, MD simulations confirmed the conformational stability of PtdIns(4,5)*P*_2_ ligand accommodated in the TH region (**Supplementary Fig. 7,8**). Thus, these results support the dynamic stability of the docked phospholipids in the TNFAIP8 hydrophobic cavity.

**Figure 4.**
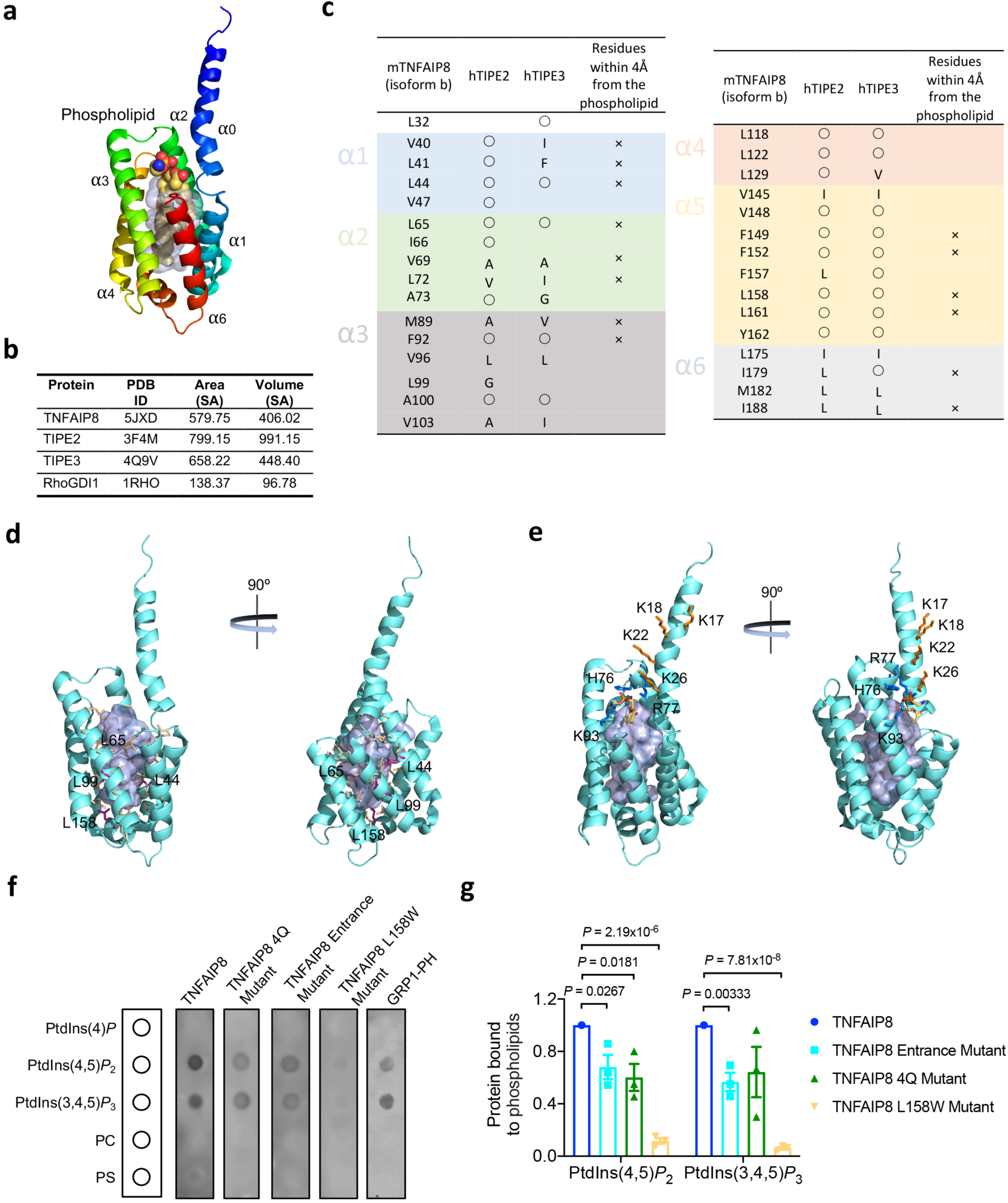
Structural analysis and mutagenesis of TNFAIP8 protein. (**a**) Cartoon presentation of murine TNFAIP8 C165S mutant (PDB accession 5JXD) in complex with a phospholipid shown by ball model. Helices are rainbow colored from *α*0 (in blue) to *α*6 (in red). The surface of the centrally located hydrophobic cavity is shown in grey. (**b**) Comparison of protein topography of murine TNFAIP8 C165S, human TIPE2, human TIPE3, and human RhoGDI1 crystal structures by CASTp 3.0. (**c**) For the 31 amino acids with hydrophobic side chains that line the cavity of murine TNFAIP8 C165S, identical (represented by circles) or similar amino acids in the corresponding positions are indicated. (**d**,**e**) Representation of mutated TNFAIP8 residues. Four conserved residues L44 (*α*1), L65 (*α*2), L99 (*α*3), and L158 (*α*5) (highlighted in magenta) out of the wall residues comprising the pocket (highlighted in yellow) were mutated to Ws to induce steric blockade in the cavity (**d**). Two positively charged residues of *α*0 helix K17 and K18 (highlighted in orange) were mutated to Qs for the 2Q mutant, and two additional residues K22 and K26 (highlighted in orange) were replaced by Qs to construct the 4Q mutant (**e**). Three positively charged amino acids H76, R77, and K93 (highlighted in blue) positioned at the opening of the cavity were mutated to Qs for the Entrance mutation (**e**). Residues are shown in stick models and cavity surface is colored grey. (**f**,**g**) Binding of 6His-SUMO tagged TNFAIP8, 4Q, Entrance, and L158W mutant proteins to the indicated lipids spotted on strips by protein-lipid overlay assay. Immunoblot was performed using antibody against 6His-tag. Signal from wild-type TNFAIP8 was set as 1. Data represent three independent experiments with similar results (**f**) or are mean ± s.e.m. of three independent experiments (**g**). *P* values are from two-sided unpaired Student’s *t*-test (**g**).

To test the importance of the hydrophobic pocket in lipid binding capacity, we compared the 31 hydrophobic amino acids comprising the pocket of murine TNFAIP8 C165S mutant to all wall residues of human TIPE2 and TIPE3 cavities and found that the amino acids in the corresponding positions were highly conserved (**Fig. 4c**). Four representative residues L44 (*α*1 helix), L65 (*α*2 helix), L99 (*α*3 helix), and L158 (*α*5 helix) at the bottom of the cavity (highlighted in magenta) out of the wall residues comprising the pocket (highlighted in yellow) were selected to mutate to the much bulkier Ws to induce steric blockade (**Fig. 4d** and **Supplementary Video 2**). To explore if the positively charged residues of *α*0 helix could contribute to phosphoinositide interactions and possibly function as a flexible lid of the cavity, we generated TNFAIP8 2Q mutant, in which K17 and K18 (highlighted in orange) of the *α*0 helix were replaced with Qs. We also mutated two additional residues K22 and K26 (highlighted in orange) in the *α*0 helix to construct the 4Q mutant (**Fig. 4e** and **Supplementary Video 3**). Furthermore, three positively charged amino acids H76, R77, and K93 (highlighted in blue) laying in the opening of the hydrophobic cavity were mutated to Qs for the Entrance mutation (**Fig. 4e** and **Supplementary Video 3**). After constructing all TNFAIP8 mutants in pET-SUMO system and introducing them to *E. coli*, we found that L44W, L65W, and L99W mutants were expressed at much lower levels judging from crude lysates when comparing to 2Q, 4Q, Entrance, or L158W mutants (data not shown), indicating that the hydrophobic cavity formed by TH domain was crucial for the structural integrity or stability of TNFAIP8 protein. TNFAIP8 4Q, Entrance, and L158W mutants were successfully purified and used for subsequent biochemical experiments (**Supplementary Fig. 9a**).

Through protein-lipid overlay assay, the binding of purified 6His-SUMO tagged TNFAIP8, 4Q, Entrance, and L158W mutant proteins to lipids on strips was determined by immunoblot using antibody against 6His-tag (**Fig. 4f**). Consistently, we found that TNFAIP8 primarily interacted with PtdIns(4,5)*P*_2_ and PtdIns(3,4,5)*P*_3_ out of the five lipids spotted. TNFAIP8 4Q and Entrance mutants exhibited marked reductions in phosphoinositide binding, while L158W mutation almost abolished this ability (**Fig. 4f,g**). For more quantitative studies, the purification of TNFAIP8 and other proteins was optimized (**Supplementary Fig. 9a**), and the size distribution of the hydrodynamics radius of molecules in solution was assessed by dynamic light scattering (DLS) (**Supplementary Fig. 9b**). A single, narrow peak at a size corresponding to the TNFAIP8 protein suggested it stable as monomers (**Supplementary Fig. 9c**). We next used surface plasmon resonance (SPR) for real-time, label-free detection of these biomolecular interactions and analysis of TNFAIP8 mutants. A supported lipid bilayer was generated on L1 sensor chip by flowing extruded small unilamellar vesicles (SUVs) containing DOPC and 3% (mole/mole) of PtdIns(4,5)*P*_2_ or PtdIns(3,4,5)*P*_3_. Sequential dilutions of TNFAIP8 or mutant proteins were injected over the surfaces while the association and disassociation were monitored (**Fig. 5a,b**). We found that murine TNFAIP8 isoform 1 protein exhibited a steady-state *K*_D_ of 4.06 ± 0.93 μM and 3.80 ± 0.20 μM to PtdIns(4,5)*P*_2_ and PtdIns(3,4,5)*P*_3_, respectively (**Fig. 5c–e**). In addition, TNFAIP8 4Q, Entrance, and L158W mutations compromised the binding affinity by 3 to 7-fold (**Fig. 5c–e**). These results indicate that *α*0 helix and TH domain are important for the ability of TNFAIP8 protein to recognize and interact with PtdIns(4,5)*P*_2_ and PtdIns(3,4,5)*P*_3_ present in lipid bilayers.

**Figure 5.**
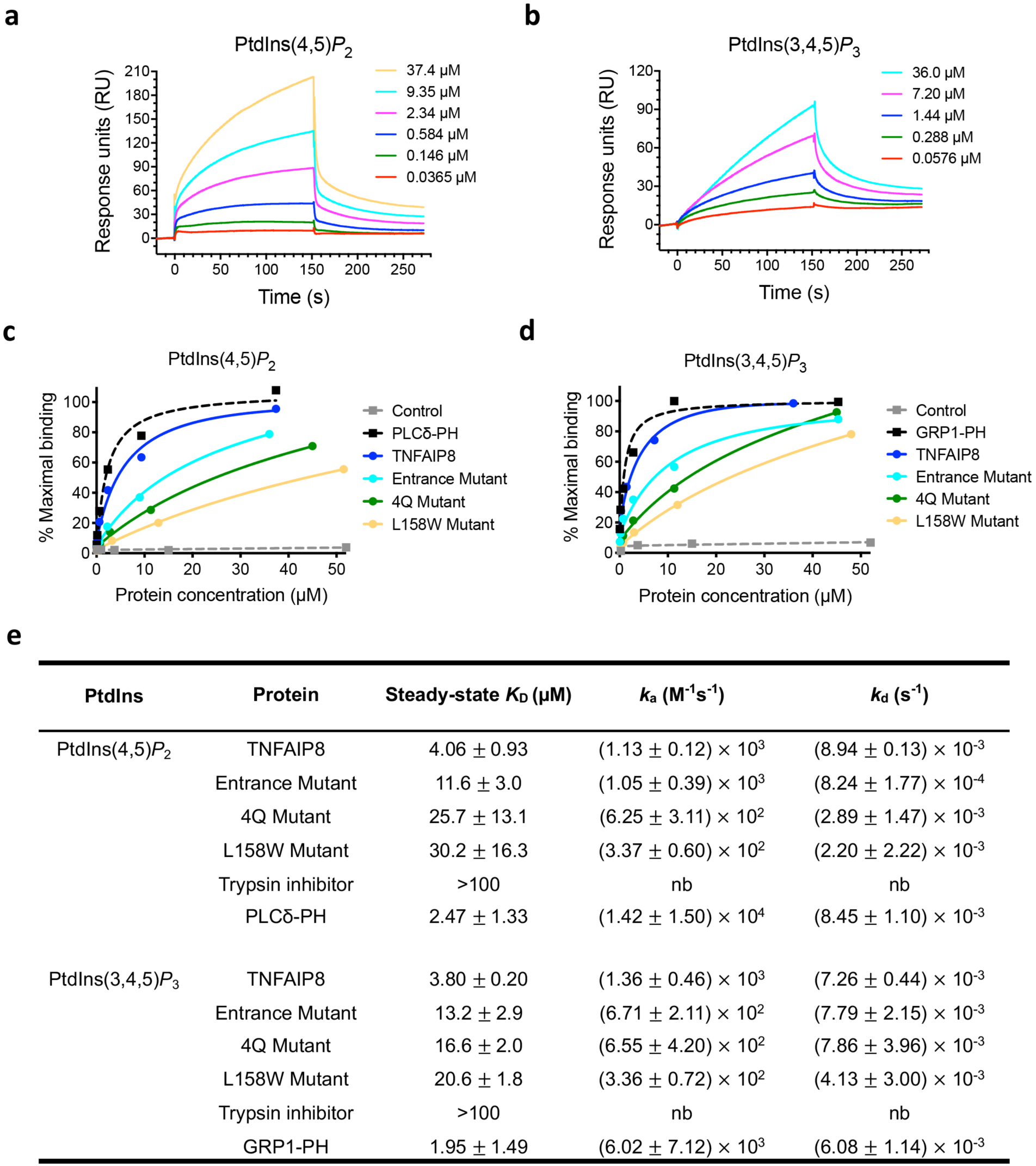
TNFAIP8 interacts with phosphoinositides through TIPE homology (TH) domain and *α*0 helix in surface plasmon resonance (SPR). (**a**,**b**) Sensorgrams monitoring the association and disassociation as sequential dilutions of murine TNFAIP8 isoform 1 were injected over the supported lipid bilayer surfaces containing DOPC and 3% (mole/mole) PtdIns(4,5)*P*_2_ (**a**) or PtdIns(3,4,5)*P*_3_ (**b**). (**c**,**d**) SPR analysis of the binding of TNFAIP8 and the indicated mutants to DOPC membranes containing 3% (mole/mole) PtdIns(4,5)*P*_2_ (**c**) or PtdIns(3,4,5)*P*_3_ (**d**) on L1 sensor chip. Purified PLCδ-PH and GRP1-PH domains were used as positive controls, and trypsin inhibitor as a negative control. (**e**) Kinetics and affinity parameters for the binding of TNFAIP8 and mutants to supported lipid bilayers containing PtdIns(4,5)*P*_2_ or PtdIns(3,4,5)*P*_3_ as measured by SPR. *k_a_*, association rate constant; *k_d_*, dissociation rate constant; *K*_D_, equilibrium dissociation constant; nb, no binding. Data represent three independent experiments with similar results (**a**–**d**) or are mean ± s.d. from three independent experiments (**e**).

### TNFAIP8 can act as a transfer protein for lipid second messengers

It has been previously demonstrated that TIPEs can function through lipid second messengers and potentiate PI3K signaling^19, 21^. Phosphoinositide extraction assay showed that when TNFAIP8 was incubated for 1 h with 100 μM vesicles containing 10% BODIPY FL PtdIns(4,5)*P*_2_ and 10% PtdIns(3,4,5)*P*_3_, increased amount of BODIPY FL PtdIns(4,5)*P*_2_ fluorescence was detected in the supernatant after ultracentrifugation, whereas equivalent experiments with trypsin inhibitor or PLCδ-PH domain showed no significant fluorescence in the supernatant (**Fig. 6a**). This result suggests that TNFAIP8 can effectively remove fluorescence labeled PtdIns(4,5)*P*_2_ from the SUVs and solubilize it in the supernatant. Additionally, the ability of extraction was compromised by the TNFAIP8 Entrance mutation (**Fig. 6b**). It is important to note that this was specific to BODIPY FL PtdIns(4,5)*P*_2_ vesicles, as neither TNFAIP8 nor Entrance mutant was able to extract BODIPY FL PC from 100 μM vesicles containing 20% BODIPY FL PC as control (**Fig. 6b**).

**Figure 6.**
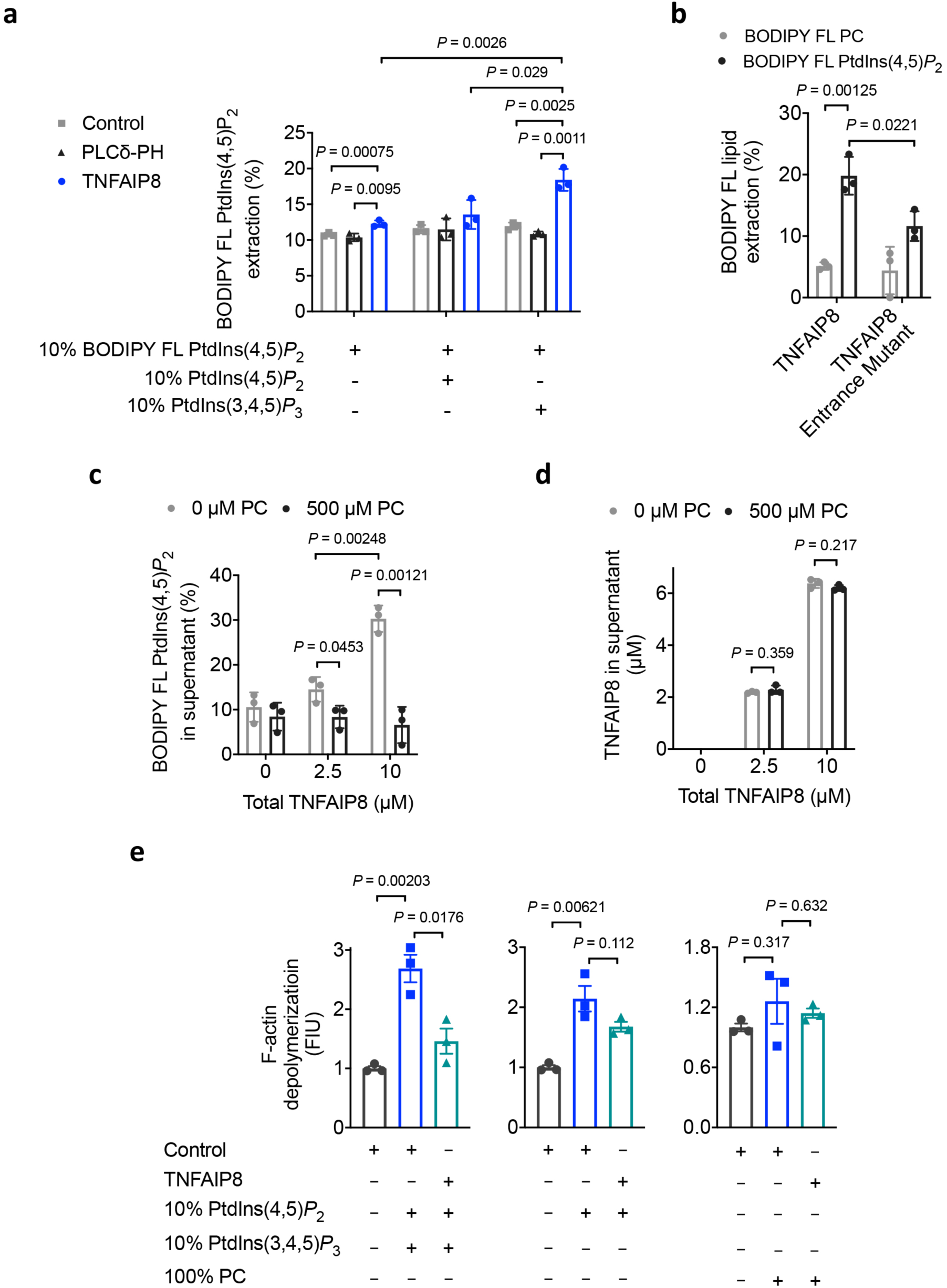
TNFAIP8 can function as a phosphoinositide transfer protein and affect cofilin-dependent depolymerization of F-actin. (**a**) Phosphoinositide extraction assay showing the proportion of PtdIns(4,5)*P*_2_ (labeled with the fluorescent dye BODIPY) extracted from small unilamellar vesicles (SUVs) containing 10% BODIPY FL PtdIns(4,5)*P*_2_ with (+) or without (-) 10% unlabeled PtdIns(4,5)*P*_2_ or PtdIns(3,4,5)*P*_3_ (grid beneath plot), in the presence of trypsin inhibitor (Control), PLCδ-PH, or TNFAIP8. (**b**) Phosphoinositide extraction assay was used to determine the percentages of BODIPY FL PtdIns(4,5)*P*_2_ extracted from 100 μM 20% BODIPY FL PtdIns(4,5)*P*_2_ vesicles and the percentages of BODIPY FL PC extracted from 100 μM 20% BODIPY FL PC vesicles in the presence of 10% unlabeled PtdIns(3,4,5)*P*_3_, and transferred to supernatant by TNFAIP8 or Entrance mutant. (**c**,**d**) The sedimentation-based transfer assay showing the depletion of BODIPY FL PtdIns(4,5)*P*_2_ from soluble TNFAIP8-BODIPY FL PtdIns(4,5)*P*_2_ in supernatant (**c**) and fractions of TNFAIP8 remaining in supernatant (**d**) after incubation with buffer alone or 100% PC vesicles. Proteins were used at a concentration of 10 μM. (**e**) Quantification of cofilin-dependent F-actin depolymerization in the presence of various combinations of control protein or TNFAIP8 plus SUVs containing 10% PtdIns(4,5)*P*_2_ with or without PtdIns(3,4,5)*P*_3_ or 100% PC (grid beneath plot). FIU, fluorescence intensity units. Data represent two (**a**–**d**) or three (**e**) independent experiments with similar results (mean ± s.d.). *P* values are from two-sided unpaired Student’s *t*-test (**a**–**e**).

To explore whether TNFAIP8 was capable of inserting extracted PtdIns(4,5)*P*_2_ into a lipid bilayer, we analyzed the transfer of BODIPY FL PtdIns(4,5)*P*_2_ from the soluble TNFAIP8/ BODIPY FL PtdIns(4,5)*P*_2_ complexes detected in the supernatant in Fig. 6a to 100% PC vesicles. We collected supernatant from the mixture of TNFAIP8 with 100 μM BODIPY FL PtdIns(4,5)*P*_2_-containing vesicles that had been subjected to ultracentrifugation, and incubated them with vesicles containing 500 μM 100% PC for 1 h. The mixture was then recentrifuged, and the level of BODIPY FL PtdIns(4,5)*P*_2_ fluorescence remaining in the supernatant was measured. Addition of 100% PC vesicles led to a substantial decrease in fluorescence (corresponding to the presumed TNFAIP8/BODIPY FL PtdIns(4,5)*P*_2_ complexes in the supernatant) (**Fig. 6c**). The concomitant loss of BODIPY FL PtdIns(4,5)*P*_2_-derived fluorescence from the supernatant was not accompanied by changes in TNFAIP8 protein concentrations in the supernatant (**Fig. 6d**), suggesting that TNFAIP8 had transferred the BODIPY FL PtdIns(4,5)*P*_2_ to the PC vesicles. These results support the hypothesis that the *α*0 helix of TNFAIP8 can function as a flexible pocket lid, and the conformational change induced by PtdIns(3,4,5)*P*_3_ binding may displace the lid (**Supplementary Fig. 5e**) and allow TNFAIP8 to extract PtdIns(4,5)*P*_2_ from the lipid bilayer, solubilize it in the aqueous phase, and transfer it to the vesicles.

### TNFAIP8 modulation of actin remodeling through phosphoinositide-binding protein cofilin

We further studied the functional significance of TNFAIP8 interaction with phosphoinositides by investigating its possible actions at the leading edge. Cofilin is a member of the actin depolymerizing factor (ADF)/cofilin family and functions as an actin-severing protein^33^. By depolymerizing older filaments, cofilin can increase the availability of filament ends and actin monomers, thereby promoting newer filamentous actin (F-actin) formation. Therefore, cofilin regulates actin dynamics by enhancing membrane protrusion and promoting cell motility and is a key regulator in the chemotaxis of metastatic cancer cells^34, 35^. The activities of cofilin are tightly controlled through several mechanisms, of which the most well-studied include phosphorylation and binding to phosphoinositides. Firstly, the severing activity of cofilin is inactivated by LIMKs or testis-specific kinases (TESKs) phosphorylation at the serine 3 (S3) position, which are regulated by Rho GTPases^33^. The dephosphorylation of S3 by slingshot homolog (SSH) or chronophin (CIN) phosphatases circumvents this inhibition. Secondly, the dephosphorylated cofilin can still be inhibited by its binding to PtdIns(4,5)*P*_2_. The local excitation global inhibition (LEGI) mechanism has been proposed for cofilin activation in polarized cells^35, 36^. This mechanism involves global phosphorylation of cytosolic cofilin resulted from external stimulation activated LIMK, and the local release and translocation of plasma membrane cofilin to the F-actin compartments (active cofilin), resulted from PLC-mediated PtdIns(4,5)*P*_2_ reduction. To understand the impact of TNFAIP8 binding to phosphoinositides on cofilin activity, we tested cofilin-mediated F-actin depolymerization in the presence or absence of purified TNFAIP8 and phosphoinositides. SUVs containing either PtdIns(4,5)*P*_2_ or PtdIns(4,5)*P*_2_ plus PtdIns(3,4,5)*P*_3_ significantly decreased cofilin-induced F-actin depolymerization (**Fig. 6e** and **Supplementary Fig. 9d–f**). TNFAIP8 could best rescue the reduction in F-actin depolymerization caused by SUVs containing both PtdIns(4,5)*P*_2_ and PtdIns(3,4,5)*P*_3_, while TNFAIP8 or control protein alone had no influence on F-actin depolymerization (**Fig. 6e** and **Supplementary Fig. 9d–f**). These results suggest that interaction of TNFAIP8 with phosphoinositides affects cofilin-induced F-actin remodeling and TNFAIP8 could mediate the activities of phosphoinositide-binding protein.

### Phosphoinositide interactions are essential for TNFAIP8-mediated cellular function

Since biochemical assays suggested that TNFAIP8 can steer cells by remodeling phosphoinositide-dependent actin dynamics, cell-based approaches were further employed to investigate the functional significances of the hydrophobic central cavity from TH domain and the electrostatic interactions of *α*0 helix. To characterize if disrupting the TNFAIP8-phosphoinositide interactions would impair cellular chemotaxis, we constructed TNFAIP8 and mutants into pGL-LU-EGFP lentiviral vector and re-expressed them in TNFAIP8-deficient HL-60 cell line (**Fig. 7a**). It was shown previously that the migration defect of dHL-60s was able to be restored by re-introducing a wild-type human TNFAIP8 isoform b transgene (**Fig. 1a,b**). Notably, the chemotaxic defect of TNFAIP8 knockout cells could be partially rescued by expressing 2Q, 4Q, or Entrance mutants, while the expression of L44W, L65W, L99W, or L158W mutants failed to upregulate the transmigration towards fMLP (**Fig. 7b**), suggesting that TNFAIP8 mutants incapable of phosphoinositide binding were incompetent to rescue the migratory defect of TNFAIP8 deficiency.

**Figure 7.**
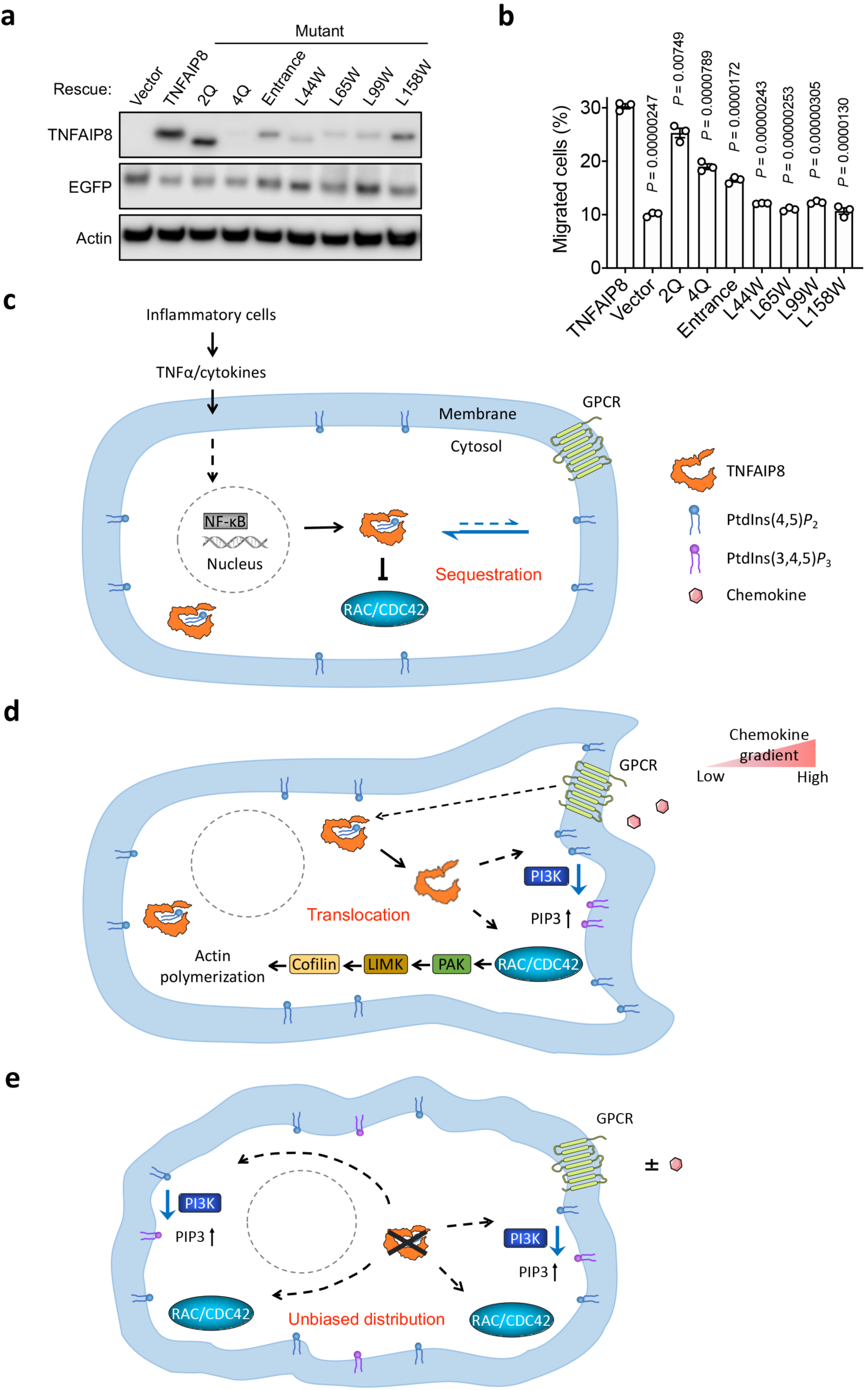
TNFAIP8 mutants deficient in phosphoinositide interactions have diminished effects on chemotaxis, and the schematic model depicting TNFAIP8 actions in cell migration. (**a**) TNFAIP8 and mutants were constructed into pGL-LU-EGFP lentiviral vectors and re-expressed in HL-60 TKO cells. Lysates were analyzed by western blot using anti-TNFAIP8 polyclonal and anti-EGFP antibodies. Actin served as a loading control. (**b**) Transwell migration assay of dHL-60 TKO cells re-expressing TNFAIP8 or mutants from lentiviral vectors (*n* = 3 biologically independent samples). Data represent two independent experiments with similar results (**a**) or three independent experiments from two clonal cell lines (mean ± s.e.m.) (**b**). *P* values are from two-sided unpaired Student’s *t*-test (**b**). (**c**) TNFAIP8 expression can be induced by inflammatory cytokines such as TNF*α* from tumor microenvironment via the activation of NF-κB. TNFAIP8 maintains the quiescent cellular state by interacting with and sequestering PtdIns(4,5)*P*_2_ and inactive Rho GTPases in the cytosolic pool. The structural basis of the inhibitory complex formation is the TNFAIP8 hydrophobic cavity for phosphoinositides and the preferential association with inactive GDP-bound form of Rac/Cdc42. This soluble pool of inactive Rac/Cdc42 acts as a reservoir in cytoplasm, allowing them to be instantly deployed to membrane for activation in response to stimuli. (**d**) GPCR activation following chemoattractant exposure generates a displacement factor or signal. This enforces plausible conformational changes of the *α*0 helix and/or hydrophobic cavity of TNFAIP8, or alternatively upregulates its membrane translocation, which can subsequently set free the Rho GTPases. The dissociated Rac/Cdc42 can then translocate to plasma membrane, where they are activated by membrane-anchored GEFs and initiate downstream effector signaling, leading to actin polymerization. The TNFAIP8 inhibitory machinery at the cell rear also supports trailing edge formation and suppresses secondary leading edge, thus conferring cues for spatial restriction and polarization maintenance. In addition, at the membrane-cytosol interface of cell front, PtdIns(4,5)*P*_2_ bound by ‘switched-on’ TNFAIP8 may serve as better water-soluble substrate for PI3Ks or influence cofilin-induced F-actin remodeling, thereby facilitating leading edge formation by enhancing phosphoinositide signaling. (**e**) TNFAIP8 knockout disrupts the plausible inhibitory complex, which results in unbiased distribution of active Rac/Cdc42 at plasma membrane and constitutively elevated PtdIns(3,4,5)*P*_3_ level, in the absence or presence of GPCR stimulation. Therefore, TNFAIP8 is built into a machinery of both short-range positive feedback and long-range inhibition, which amplifies the shallow chemical gradients into axes of cell polarity and is essential for expanding the range of chemotaxic sensitivity.

## DISCUSSION

TNFAIP8 expression has been reported to be induced by TNF*α* and high glucose stimulation, as well as activation of NF-κB in multiple cell types^37, 38^. The orphan nuclear receptor chicken ovalbumin upstream promoter transcription factor I (COUP-TFI) is identified as a transcriptional repressor of TNFAIP8 gene^39^, which is involved in the induction of TNFAIP8 promoter by TNF*α*. However, the detailed transcriptional control mechanisms of TNFAIP8 remain elusive. Current evidences support the notion that TNFAIP8 links a molecular bridge from inflammation to cancer by combining TNF*α*/NF-κB pathway to phosphoinositide signaling (**Fig. 7c**). We observe that TNFAIP8-deficient cells have reduced migratory and adhesive capacity, upregulated PtdIns(3,4,5)*P*_3_ level, as well as aberrant molecular pathways downstream of Rac/Cdc42 activation. These results imply that TNFAIP8 can maintain the quiescent cellular state by interacting with and sequestering PtdIns(4,5)*P*_2_ and inactive Rho GTPases in the cytosolic pool (**Fig. 7c**). Since TIPE2 preferentially associates with dominant negative Cdc42-17N but not constitutively active Cdc42-61L mutant, it is possible that TIPEs stabilize the plausible inhibitory complex by associating with the GDP-bound form of Rac/Cdc42. This soluble pool of inactive Rac/Cdc42 acts as a reservoir in cytoplasm, allowing them to be instantly deployed to membrane for activation in response to stimuli. Following chemoattractant exposure, TNFAIP8 can steer cells along the gradient by regulating Rho GTPases and phosphoinositide-binding proteins through two distinct mechanisms (**Fig. 7d**). Firstly, GPCR activation generates a displacement factor or signal; this signal may enforce conformational changes of TNFAIP8 (the flexible *α*0 helix is likely to act as a lid and switch the configuration of hydrophobic cavity) or alternatively upregulate its membrane translocation, which can subsequently set free the Rho GTPases. Mechanisms for the formation and release of cytosolic TNFAIP8-Rho GTPase complex remain unclear, although protein-protein interaction or phosphorylation might play a role in the stability of the inhibitory complex. The dissociated Rac/Cdc42 can then translocate to plasma membrane, where they are activated by membrane-anchored GEFs and initiate downstream effector signaling through the PAK-LIMK-cofilin pathway, leading to actin polymerization. In addition, TNFAIP8 abundance may expedite both the delivery and partition of Rho GTPases, contributing to fast signal termination. The TNFAIP8 inhibitory machinery at the cell rear also supports trailing edge formation and suppresses secondary leading edge, thus conferring cues for spatial restriction and polarization maintenance. Secondly, at the membrane-cytosol interface of cell front, the ‘switched-on’ TNFAIP8 can influence the activities of phosphoinositide-binding proteins to facilitate leading edge formation. For instance, PtdIns(4,5)*P*_2_ with fatty acid chains accommodated in the hydrophobic pocket may serve as better water-soluble substrate for PI3Ks, and TNFAIP8 interaction with PtdIns(4,5)*P*_2_ could enhance cofilin-induced F-actin remodeling. Therefore, TNFAIP8 is built into a machinery of both short-range positive feedback and long-range inhibition, which amplifies and translates the shallow chemical gradients into axes of cell polarity (**Fig. 7d**). Consequently, TNFAIP8 knockout disrupts the plausible multimolecular inhibitory complex, which results in unbiased distribution of active Rac/Cdc42 at plasma membrane and constitutively elevated PtdIns(3,4,5)*P*_3_ level, in the absence or presence of GPCR stimulation (**Fig. 7e**).

Rho family of small GTPases are well known to be regulated by GEFs, GAPs, and RhoGDIs. RhoGDIs can inhibit Rac by extracting prenylated Rac-GDP from plasma membrane and by decreasing the rate of nucleotide dissociation. RhoGDIs have also been reported to be able to prevent the degradation of Rho GTPases. Our study reported here indicates that TIPE family might represent a unique class of negative regulators of Rho GTPases, distinct from the aforementioned molecules. Due to the potential functional overlap of both classes of Rho GTPase inhibitors, double deficiency of TNFAIP8 and RhoGDIs would be likely to generate more dramatic abnormalities in migratory cells or model animals. Even though TNFAIP8 assumes some of the properties of RhoGDIs by sequestering inactive Rho GTPases, it might mediate more selective release and activation of a single Rho GTPase, given that each mammalian RhoGDI interacts with multiple Rho GTPases. The interplay between GEFs, GAPs, RhoGDIs, and TIPE proteins and their coordinate regulation need to be studied in a combined manner.

Numerous phosphoinositide-binding domains have been discovered so far, including PH, PX, PHD, FYVE, FERM, and ENTH domains^40^. In contrast to the canonical binding modes of these domains that target phosphoinositide head groups, TH domain accommodates the lipid tails. It would be interesting to further visualize the subcellular changes of phosphoinositide levels regulated by TH domain expression and monitor the alternations upon stimulations. TH domain is conserved through evolution and is found in invertebrates such as fruit fly, unicellular eukaryotes such as Entamoeba, as well as vertebrates. Interestingly, G protein interacting protein 1 (Gip1) has recently been found to regulate GPCR signaling of chemotaxis in eukaryote Dictyostelium discoideum by binding and sequestering heterotrimeric G proteins in the cytosolic pool. Remarkably, the crystal structure of the C-terminal region of Gip1 showed cylinder-like folding with a central hydrophobic cavity, in homology to TIPE TH domain^41, 42^. Dictyostelium deficient in Gip1 exhibited severe defects in following the gradient of the chemoattractant cyclic adenosine monophosphate (cAMP), indicating that Gip1 was essential in regulating the directional migration of Dictyostelium discoideum^41^. These findings suggest that TIPEs, along with Gip1, may constitute a novel family of evolutionarily conserved proteins with common tertiary structure features to direct eukaryotic cell chemotaxis. In addition, TNFAIP8 was also reported to directly bind to G*α*i and couple with G*α*i-coupled dopamine-D2short (D2S) receptor to reduce cell death and induce cellular transformation^43^. Through yeast two-hybrid screening and co-immunoprecipitation or pull-down confirmation, TNFAIP8 was shown to preferentially interact with G*α*i, but not other G*α* proteins that not associated with chemokine receptors. Deletion of the last 15 amino acids of TNFAIP8 C-terminus was sufficient to disrupt this interaction^43^. Thus, it is likely that TNFAIP8 senses chemokine gradient through GPCRs and further study will clarify if G*α*i proteins are responsible for recruiting TNFAIP8 to the activation site.

The functions of TIPE family may also be differentially regulated by other sequences present in the proteins, such as the N-terminal regions. For example, the unique N-terminus of TIPE3, which is not shared by other family members (**Supplementary Fig. 1a**), has been reported to act in a dominant negative manner^21^. This raises the question as to whether the many transcript variants and protein isoforms of TNFAIP8 in both mice and human may be transcriptionally regulated to exert diverse cell-type specific functions. Our results indicate that TNFAIP8 can play a role in the modulation of PI3K-AKT-mTOR and MEK-ERK pathways, and some studies suggest that it may be an inhibitor of caspases-induced apoptosis^24, 44^. More work will be needed to elucidate the roles of TNFAIP8 in cell proliferation and death signaling.

A better understanding of cancer cell motility and relevant mechanisms could be useful in developing therapeutic strategies to abate metastasis-initiating migration in the clinic to combat metastasis. Our study reveals that TNFAIP8, as a novel negative regulator of Rho GTPases, expands the range of chemotaxic sensitivity in cancer cells, and advises effective pharmaceutical drug design targeting TNFAIP8 for its phosphoinositide interaction capacity.

## ACKNOWLEDGEMENTS

We thank Erfei Bi (University of Pennsylvania) for the conceptualization of TNFAIP8 inhibitory theory; Tatyana Svitkina, Paula Oliver, and Wei Guo (University of Pennsylvania) for valuable discussions and support. We thank Xinyuan Li, Ali Zamani, Amanda Boggs, and Chen lab members for technical support and reagents. We also thank the Molecular Screening Core (The Wistar Institute) and Biophysical and Structural Biology Core (University of Pennsylvania) for technical assistance. This work was funded by the National Institutes of Health, USA (R01-AI099216, R01-AI121166, R01-AI136945, and R56-AI-132329 to Y.H.C.).

## AUTHOR CONTRIBUTIONS

M.L. and Y.H.C. conceived the study; M.L. designed, executed, and analyzed the experiments; H.S. oversaw animal husbandry, processed murine tissues, and analyzed the data; S.A.F. designed certain biochemical experiments and helped with data analysis; P.H. performed TCGA survival analysis; M.L. wrote the manuscript with input and comments from all authors; Y.H.C. supervised the study.

## MATERIALS & CORRESPONDENCE

Methods, including statements of data availability and any associated accession codes and references, are available in the online version of this paper. Note: Supplementary Information is available in the online version of the paper.

## DATA AVAILABILITY

Data supporting the findings of this manuscript are available from the corresponding authors upon request.

## METHODS

### Cell culture

The cell lines were obtained from the American Type Culture Collection (ATCC) or Clontech. HEK293T, Lenti-X 293T, PANC-1 pancreatic carcinoma, EL-4 T cell lymphoma, WEHI 7.1 T cell lymphoma, Hepa 1-6 hepatoma, B16-F0 melanoma, B16-F10 melanoma, Fibrosarcoma 573, and Raw 264.7 macrophage cell lines were cultured in Dulbecco’s Modified Eagle Medium (DMEM) with 10% FBS, 1% L-glutamine, and 1% Penicillin-Streptomycin (D10 medium). U-937 lymphoma, NCI-H69 small cell lung cancer, NCI-H727 lung cancer, AsPC-1 pancreatic adenocarcinoma, and MT-901 mammary carcinoma cell lines were cultured in RPMI-1640 with 10% FBS, 1% L-glutamine, and 1% Penicillin-Streptomycin (R10 medium). T24 urinary bladder cancer and HT-29 colorectal adenocarcinoma cell lines were cultured in McCoy’s 5A medium containing 10% FBS and 1% Penicillin-Streptomycin. HEC-1-B uterus adenocarcinoma and Neuro-2a neuroblastoma cell lines were cultured in Eagle’s Minimum Essential Medium (EMEM) composed of Minimum Essential Medium (MEM) containing 1× Non-Essential Amino Acids (NEAA), 1% L-glutamine, 1 mM sodium pyruvate, 10% FBS, and 1% Penicillin-Streptomycin. SW480 colorectal adenocarcinoma cell line was cultured in Leibovitz’s L-15 medium containing 10% FBS and 1% Penicillin-Streptomycin. HL-60 promyelocytic leukemia cell line was cultured in R10 medium supplied with 25 mM HEPES. HL-60 cell concentration was kept between 1.5×10^5^ to 10×10^5^ cells/ml during passage. Frozen cells of the same lot were thawed every two months for the same quality. HL-60 cells were differentiated by 1.25% (vol/vol) DMSO and 5 ng/ml human G-CSF continuously in culture for 4–6 days before harvest for experiment (cell density was kept at 1.5–2.5×10^5^ cells/ml). Cell cultures were examined periodically for bacteria contamination and tested for mycoplasma by LookOut Mycoplasma PCR Detection Kit (Sigma).

### Constructs and plasmid transfection

Expression plasmids of human TNFAIP8 isoform b, murine TNFAIP8 isoform 1, and the PH domains from phospholipase C-δ1 (PLCδ-PH) and general receptor for phosphoinositides (GRP1-PH) were constructed by cloning PCR-amplified cDNA into pET-SUMO vector (LifeSensors), in frame with the N-terminal 6His-SUMO tag. The mutagenesis was performed using Stratagene QuikChange II Mutagenesis Kit (Agilent) by following the manufacturer’s protocol. The lentiviral expression plasmids of TNFAIP8 and mutants were constructed by cloning the open reading frame of each cDNA into the multiple cloning site of pGL-LU-EGFP vector. Full-length human TNFAIP8 isoform b or murine TIPE2 cDNA was cloned in frame with a N-terminal GST tag into pRK5 expression vector. Wild-type Cdc42 cDNA was obtained from Addgene and subcloned into pRK5 with a Myc tag at the N-terminus. Cdc42-17N and Cdc42-61L were generated by mutagenesis from wild-type Cdc42. HEK293T cells were transfected with plasmid DNA using Fugene 6 reagent (Promega) by following the manufacturer’s protocol.

### Generation of TNFAIP8-deficient single clonal cell line by CRISPR/Cas9

TNFAIP8-deficient HL-60 cells were generated using CRISPR/Cas9 by TNFAIP8-specific single guide RNA (sgRNA) or non-targeting control single guide RNA (sgControl) as described^45^. Three sgRNAs were designed to target the earliest sequences of the shared exon of human TNFAIP8 isoforms with optimal on-target and off-target scores (Supplementary Table 1). CRISPR target sequences were cloned into LentiCRISPRv2 backbone by adding BsmBI digestion sites using specific primers (Supplementary Table 2). The constructs were transfected into Lenti-X 293T cells, and the harvested lentiviruses were concentrated and titrated. With a multiplicity of infection (MOI) of 10 and functional lentivirus titer of 1.2×10^6^ colony forming unit (CFU)/ml, the transduction efficiency of HL-60 cells measured after 48 h was over 80%. The medium was replaced with selection medium containing 1 μg/ml puromycin at 48 h post-transduction as well as every 2–3 days until antibiotic-resistant colonies could be identified (around 4–7 days). T7EI genomic cleavage detection (GCD) assay (IDT) was performed on the genomic DNA of cells harvested before or three days after puromycin selection. Specific primers amplifying the targeted sequences (Supplementary Table 2) were used to confirm the efficiency of nucleotide insertion or deletion (indel) formation. The puromycin-resistant cells were passed through a 40 μm cell strainer, and this single cell suspension was plated at a suitable density to TCS semi-solid methylcellulose-based medium (ClonaCell) by following the manufacturer’s protocol. After 10–14 days the cultures were inspected for the presence of colonies, and the single clones were picked and expanded in liquid medium for cryopreservation and further analysis.

### Lentivirus infection

Lenti-X 293T cells were pre-cultured in lentivirus packaging medium containing Opti-MEM I reduced serum medium, 1× GlutaMAX, 1 mM sodium pyruvate, and 5% FBS. Expression constructs were transfected into Lenti-X 293T using Lipofectamine 3000 reagent (Thermo Fisher Scientific) by following the manufacturer’s protocol. Lentivirus was produced by co-transfecting constructed expression vectors along with 3^rd^ generation lentiviral packaging plasmids (pRSV-Rev, pCMV-VSV-G, and pMDLg/pRRE). Two batches of lentivirus-containing media were harvested 24 h and 52 h after transfection, followed by filtering through a 0.45 μm Steriflip PVDF membrane (MilliporeSigma). Lentivirus was concentrated 100-fold using Lenti-X Concentrator (Clontech) and titrated by RT-qPCR Titration Kit (Clontech) as well as EGFP-positive cell counts. Transduction was performed by incubating cells with medium containing the lentivirus at a MOI of 10–100 in the presence of 4 μg/ml Polybrene (Sigma) for 12–24 h. The amount of lentivirus used to re-express target protein in knockout cells was titrated and adjusted to ensure close to the endogenous level.

### Transwell migration and invasion assays

The migration of dHL-60s towards chemoattractants was performed using 3 μm pore Transwell filters (Corning) as described^46^. dHL-60 cells were washed two times and rested in migration assay buffer for 30 min at 37°C. Migration assay buffer was composed of Hanks’ Balanced Salt Solution (HBSS) with Ca^2+^/Mg^2+^, 25 mM HEPES, and 0.2% freshly dissolved, fatty acid and endotoxin free human serum albumin (HSA, Sigma). For the invasion assay, filters were coated with 50 μg/ml Matrigel (Corning) and placed at room temperature for 1 h to form a thin gel layer. 2×10^5^ cells were added using reverse pipetting method to avoid bubbles. When studying the effects of inhibitors on transmigration, cell seeding volume was reduced to half to allow for the addition of an equal volume of test reagents in 2× concentration, and cells were pre-incubated with the antagonists for 45 min at 37°C. The inhibitors were used at the following final concentrations: LY-294,002 (100 μM, Sigma), NSC 23766 (300 μM, Sigma), Torin1 (250 nM, Calbiochem), InSolution Z62954982 (100 μM, Calbiochem), Y-27632 (10 μM, Cell Signaling Technology), TH-257 (5 μM, Sigma), and FRAX486 (10 μM, Sigma). fMLP or CXCL8 was added to the bottom wells at a concentration of 10 nM or 3 nM, respectively. dHL-60 cells were allowed to migrate for 1.5 h at 37°C. Pre-warmed cell detachment solution (0.05% Trypsin-EDTA without phenol red) was added to the lower chamber to dissolve any cells attached to the bottom of the insert. The numbers of cells that migrated into the lower chamber as well as those detached by Trypsin-EDTA were quantified by CellTiter-Glo (CTG) assay (Promega). % Migration = the number of cells that migrated in response to chemoattractant / the total number of cells seeded on the chamber.

### Static adhesion assay

The 96-well cell culture plates were coated with 50 μg/ml fibronectin or 5 μg/ml ICAM-1 for 2 h at 37°C. dHL-60 cells were serum starved for 2 h in assay buffer containing HBSS with Ca^2+^/Mg^2+^, 25 mM HEPES, and 0.2% HSA. Cells were subsequently stimulated with fMLP (100–200 nM) or TNFα (30 ng/ml) and seeded on fibronectin or ICAM-1 coated surfaces at 1.5×10^5^ cells per well. After incubation at 37°C for 30 min, the plates were gently washed three times with HBSS buffer (containing Ca^2+^/Mg^2+^) to remove cells that were not able to form tight contacts with the ligands. Cells in HSA-coated negative control wells should be rare under microscope; if not, washing was repeated. The cells left in the wells were quantified by CTG assay (Promega), and the average percentages of adherent cells were calculated relative to unwashed wells that represented 100%.

### Western blot and subcellular fractionation

Cells were lysed for 15 min at 4°C in RIPA lysis buffer (Sigma) containing 50 mM Tris-HCl (pH 8.0), 150 mM sodium chloride, 1.0% Igepal CA-630, 0.5% sodium deoxycholate, 0.1% sodium dodecyl sulfate (SDS), cOmplete protease inhibitor cocktail (Roche), and PhosSTOP phosphatase inhibitors (Roche). Cell debris was removed by centrifugation at 14,000g for 15 min at 4°C. For subcellular fractionation, Qproteome Cell Compartment Kit (Qiagen) was used to separate membrane, cytoplasmic, nuclear, and cytoskeleton proteins by following the manufacturer’s protocol. Protein concentration of lysate was measured by BCA Protein Assay (Pierce). Equal amount of total protein was separated by SDS–PAGE, transferred to PVDF membranes, immunoblotted with specific primary and secondary antibodies, and the signals were revealed by chemiluminescence (Pierce). Anti-GST, GST-HRP, 6His-HRP, Myc, AKT, cofilin, p-cofilin (S3), p-LIMK1 (T508)/LIMK2 (T505), p-p38 (T180/Y182), p-Erk1/2 (T202/Y204), p-p70 S6 Kinase (T389), p-p70 S6 Kinase (S371), p-4E-BP1 (T37/46), cyclin D1, p-PAK1 (T423)/PAK2 (T402), PAK1/2/3, ATPase, GAPDH, integrin-β1, and Histone H3 antibodies were purchased from Cell Signaling Technology. Anti-TNFAIP8 and TIPE2 antibodies were purchased from Proteintech. Anti-Par3, EGFP, and actin (monoclonal, AC-15) antibodies were purchased from Sigma-Aldrich. Control IgG was purchased from Santa Cruz Biotechnology. Anti-rabbit IgG-HRP (NA934) and anti-mouse IgG-HRP (NA931) were purchased from GE Healthcare Life Sciences. The densitometric quantification of western blot signals was performed using ImageStudioLite (LI-COR Biosciences) and ImageJ softwares.

### Survival analysis

Available patient survival data were obtained from TCGA project, and patients were ranked based on their TNFAIP8 expression. The lower half of ranked patients was defined as the ‘bottom 50% expression’ group and the top half was defined as the ‘top 50% expression’ group. Kaplan–Meier survival analysis and log-rank significance test were performed accordingly on these two patient groups.

### Lipid extraction

Total lipids were isolated from cells as described^47^, and stock solutions were prepared using standard glassware. Cells were pelleted, washed, and resuspended in HBSS buffer containing Ca^2+^/Mg^2+^ at 3×10^7^ cells/ml. Aliquots of 20 μl cells were saved to harvest actin protein as loading control. The left cells were divided equally into two groups in 15 ml conical tubes and were stimulated with 10 nM fMLP or DMSO vehicle for 60 s at 37°C. Reactions were terminated by addition of 1500 μl quench mix (75 ml stock solution was comprised of 48.4 ml methanol, 24.2 ml chloroform, and 2.355 ml 1 M HCl). The above steps created primary lipid extracts from cells, which resulted in a thoroughly mixed, single-phase sample. 1450 μl chloroform was then added, followed by 30 s vortex and addition of 340 μl 2 M HCl. The sample was then thoroughly mixed again and centrifuged at 1500g for 5 min at room temperature. This created a two-phase system with an upper aqueous layer and a lower organic phase, with a protein band at the interface. The upper phase was siphoned, and the lower organic phase was collected into a 2 ml safe-lock polypropylene tube. The extracted lipids were dried down with light nitrogen stream, and then speed-vacuumed at 35°C for 1 h. The dried lipids were stored at −20°C or −80°C and dissolved in a 1:1 mixture of chloroform and methanol containing 0.1% HCl before experiments.

### Protein-lipid overlay assay

Cellular levels of PtdIns(3,4,5)*P*_3_, PtdIns(4,5)*P*_2_, or PtdIns(4)*P* were measured by protein-lipid overlay with GST-GRP1-PH, GST-PLCδ-PH, or GST-SidC-3C domain, respectively. Aliquots of 1–2 μl extracted lipids were spotted onto nitrocellulose membrane and allowed to dry in dark at room temperature for 1 h or at 4°C overnight. The membranes were then blocked with TBS-T (10 mM pH 8.0 Tris-HCl, 150 mM NaCl, and 0.05% Tween 20) containing 3% fatty acid-free BSA (Sigma) and gently agitated for 1 h at room temperature. The membranes were subsequently incubated with 0.5–1.0 μg/ml GST-PLCδ-PH, GST-GRP1-PH, or GST-SidC-3C (Echelon) in TBS-T with 3% fatty acid-free BSA for 1 h at room temperature. Proteins were detected by anti-GST-HRP antibody (Cell Signaling Technology) and chemiluminescence (Pierce). Control membranes in dot blots were stained with anti-GST-HRP only. For normalization, a small aliquot of cells was lysed in RIPA lysis buffer with protease and phosphatase inhibitors. The lysates were spotted on membrane and immunoblotted with anti-actin (1:4000 dilution) and HRP-conjugated secondary anti-mouse IgG (1:3000 dilution). For the protein-lipid overlay of 6His-SUMO tagged TNFAIP8 and mutants, synthetic lyophilized lipids were reconstituted to 1 mM stocks in a 1:1 solution of methanol and chloroform and stored at −80°C before use. DOPC (Avanti Polar Lipids), DOPS (Avanti Polar Lipids), PtdIns(4)*P* (CellSignals), PtdIns(4,5)*P*_2_ (CellSignals), and PtdIns(3,4,5)*P*_3_ (CellSignals) were spotted onto nitrocellulose membranes and processed as described above. The membranes were then incubated with 10 nM purified TNFAIP8 or mutant proteins in TBS-T with 3% fatty acid-free BSA, and the binding was detected with anti-6His-HRP antibody (Cell Signaling Technology). Signals were revealed by chemiluminescence (Pierce) and quantified by desitometry using ImageStudioLite software (LI-COR Biosciences).

### GST pull-down assay

Cells were transfected with GST-tagged ‘bait’ protein and ‘prey’ protein expressing plasmids, lysed, and quantified by BCA assay to adjust to 1–5 mg/ml. The lysates with equal total protein amount were incubated with 50 μl pre-washed Glutathione Sepharose 4B (GE Healthcare Life Sciences) in 100 μl GST pull-down buffer (20 mM Tris, 150 mM NaCl, 2 mM MgCl2, 0.1% NP-40, and 20 μg/ml BSA) for 2 h at 4°C with gentle rotation. Protein-bound Sepharose was washed 3 times, eluted with glutathione, and boiled in 2× Laemmli sample buffer. Supernatants were subjected to SDS–PAGE and immunoblot analysis.

### Co-immunoprecipitation assay

Immunoprecipitation was performed using Dynabeads protein G (Invitrogen) by following the manufacturer’s protocol. Briefly, 1.5 mg protein G Dynabeads were coated with 5 μg specific antibodies or IgG control for 1 h at room temperature with rotation. After removing unbound antibody, the bead-antibody complex was incubated with 500 μl cell lysates for 4 h at 4°C with gentle rotation. The captured Dynabead-Ab-Ag complex was washed four times with PBS and boiled in 2× Laemmli sample buffer. The eluted proteins were fractionated by SDS–PAGE and detected by western blot.

### Mice

*Tnfaip8^-/-^* and *Tipe2^-/-^* C57BL/6 mice were produced as previously described^19^. The *Tnfaip8^-/-^ Tipe2^-/-^* double-knockout (DKO) mice were generated by crossing *Tnfaip8^-/-^* with *Tipe2^-/-^* mice. WT C57BL/6 mice were purchased from Jackson Laboratories. Mice were housed in the Animal Care Facilities of University of Pennsylvania under pathogen-free conditions. Animal procedures were all pre-approved by the Institutional Animal Care and Use Committee of the University of Pennsylvania. All mice experiments conformed to the relevant regulatory standards.

### Molecular docking

Crystal structure of TNFAIP8 C165S mutant (5JXD) with a resolution of 2.03 Å was retrieved from PDB. The 3D structure was refined and optimized by the protein preparation wizard from Schrödinger Maestro software, including adding hydrogen atoms, removing water molecules, and fixing the bonds and orientations^48^. The structures of phospholipids were prepared by Schrödinger LigPrep, which optimized the geometry and generated 3D molecules with minimized energy. OPLS 2005 force field was used to model protein-ligand interactions. The molecular docking was performed by following the Schrödinger Glide protocol. Extra precision (XP) sampling was employed, and the parameters were set to the default values of the software. The molecular mechanics combined with generalized Born and surface area (MM-GBSA) scoring function was used to estimate binding free energies by following the Schrödinger Prime protocol. All molecular graphics were produced with Schrödinger PyMOL Molecular Graphics System (Version 2.3).

### Molecular dynamics (MD) simulations

MD simulations were carried out using Schrödinger Desmond package^49^. Each docked protein-ligand complex was surrounded with a cubic box sized 10 × 10 × 10 Å. Water molecules were explicitly described by the simple point charge (SPC) model. A salt concentration of 0.15 M and OPLS 2005 force field parameters were used in all simulations. Energy minimization was performed, and the simulation system was equilibrated by following Desmond’s default values. The simulation length was 50 ns with a relaxation time of 1.0 ps, and the temperature of 300 K and 1.0 bar pressure were applied in all runs. The Martyna–Tuckerman–Klein chain coupling scheme was used for the pressure control with a constant of 2.0 ps, and the Nosé–Hoover chain coupling scheme was used for the temperature control with a constant of 1.0 ps. The long-range electrostatic interactions were evaluated using the particle mesh Ewald method (PME). The cutoff radius in coulombic interactions was 9.0 Å. Non-bonded forces were calculated using the RESPA integrator with an outer time step of 6.0 fs and inner time step of 2.0 fs. The stability of MD simulation was monitored by checking the root mean square deviation (RMSD) of the ligand and protein atom positions in time, and the interactions were visualized by the Desmond Simulation Interaction Diagram tool.

### Protein purification and dynamic light scattering (DLS)

Constructs encoding the 6His-SUMO tagged proteins were transformed into chemically competent BL21(DE3) *Escherichia coli* (Agilent). A fresh bacteria colony was inoculated to 10 ml LB broth containing 200 μg/ml Ampicillin and 50 μg/ml Chloramphenicol and grown at 37°C overnight. The seed culture was 1:50 diluted to Terrific Broth containing antibiotics and 0.2% glucose, and the cells were grown at 37°C until an OD600 of 0.6 was reached. Protein expression was induced by adding 0.5 mM IPTG and rocking vigorously at 16°C overnight. After centrifugation, collected bacterial pellets were resuspended in lysis buffer supplemented with 1 mg/ml lysozyme. The lysis buffer was composed of 50 mM sodium phosphate, 300 mM NaCl, 10% glycerol, 10 mM Imidazole, and 0.1% (w/v) CHAPS, and adjusted to pH 7.6. 20 mM 2-Mercaptoethanol and cOmplete EDTA-free protease inhibitors (Roche) were added to the pre-cold lysis buffer right before use. After incubation on ice for 30 min, cells were disrupted by sonication using 6×10 s bursts at 200–300W with a 10 s cooling period between each burst. Benzonase nuclease (Sigma) was added at the amount of 3 units per ml of original culture volume processed. Cleared lysates after centrifugation were added to equilibrated Ni-NTA Agarose (Qiagen) and mixed gently on a rotary shaker at 4°C for 60 min. The Ni-NTA matrix was washed 4 times by spinning down at 1000g for 1 min and discarding the supernatant. The wash buffer was composed of 50 mM sodium phosphate, 500 mM NaCl, 10% glycerol, and 30 mM Imidazole, and adjusted to pH 7.6. The protein was eluted 4–5 times with 1 ml elution buffer by gently rotating for 2 min at 4°C. The elution buffer was composed of 50 mM sodium phosphate, 500 mM NaCl, 10% glycerol, 250 mM Imidazole, and 0.1% CHAPS, and adjusted to pH 7.6. The eluates were exchanged to SUMO protease buffer using Slide-A-Lyzer cassettes (Thermo Fisher Scientific) at 4°C overnight, which was coupled with the cleavage step with SUMO Protease 1 added. The SUMO protease buffer was composed of 50 mM sodium phosphate, 300 mM NaCl, 10% glycerol, 0.01% CHAPS, and 1 mM DTT, and adjusted to pH 7.2. The cut 6His-SUMO tag, uncut 6His-SUMO fusion protein, and 6His-tagged SUMO Protease 1 in the post-cleavage solution were removed by a second round of Ni-NTA affinity chromatography. The amount of Ni-NTA agarose required to capture the 6His-tagged contaminants was calculated. Ni-NTA resin was equilibrated with SUMO protease buffer and mixed gently at 4°C for 10 min with dialyzed protein solution. Untagged native proteins were eluted, separated by SDS–PAGE, and were at least 95% pure judging from overloaded Coomassie Blue G-250 stained gels. Protein concentrations were determined based on absorbance at 280 nm using calculated extinction coefficients. For long-term storage, proteins were dialyzed against 1 L storage buffer, which was composed of 50 mM Tris-HCl, 50 mM NaCl, and 25% glycerol. The storage buffer was supplemented with 0.02% (w/v) sodium azide, 5 mM TCEP, and 1 mM EDTA. Single-use amount was aliquoted to sterile tubes, snap frozen, and stored in −80°C freezer. After thawing and exhaustive dialysis in HBS buffer, the size distribution of hydrodynamics radius for molecules in solution was measured by DynaPro NanoStar (Wyatt Technology) following the manufacturer’s protocol.

### Phosphoinositide extraction and transfer assays

The small unilamellar vesicles (SUVs) were produced and sedimentation-based phosphoinositide extraction and transfer assays were performed as previously described^18, 19^. Dioleoylphosphatidylcholine (DOPC) and brominated distearoyl PC (brominated PC) were purchased from Avanti Polar Lipids. PtdIns(4,5)*P*_2_ and PtdIns(3,4,5)*P*_3_ were purchased from CellSignals. BODIPY FL PC and PtdIns(4,5)*P*_2_ were purchased from Echelon Biosciences. Purified trypsin inhibitor of *Glycine max* was purchased from Sigma-Aldrich and used as a control protein. For the sedimentation-based PtdIns(4,5)*P*_2_ extraction assay, 100 μM SUVs composed of 10% BODIPY FL PtdIns(4,5)*P*_2_ in DOPC background with or without 10% unlabeled PtdIns(4,5)*P*_2_ or PtdIns(3,4,5)*P*_3_ were mixed with 10 μM purified trypsin inhibitor, PLCδ-PH, or TNFAIP8, incubated for 1 h at room temperature, and subjected to ultracentrifugation as described^50^. In sedimentation-based PC extraction assay, 100 μM SUVs composed of 10% BODIPY FL PC with 10% unlabeled PtdIns(3,4,5)*P*_3_ were mixed with 10 μM purified TNFAIP8 or Entrance mutant, incubated for 1 h at room temperature, and centrifuged as described above. The fluorescence intensity of supernatant (corresponding to BODIPY FL PtdIns(4,5)*P*_2_ or BODIPY FL PC extraction) was detected using the Infinite 200 Pro fluorescence plate reader (Tecan). In sedimentation-based PtdIns(4,5)*P*_2_ transfer assay, supernatant generated in PtdIns(4,5)*P*_2_ extraction assay was mixed with 500 μM 100% PC SUVs or HBS buffer alone, incubated for 1 h at room temperature and centrifuged. The fluorescence intensity of supernatant (corresponding to soluble TNFAIP8-BODIPY FL PtdIns(4,5)*P*_2_) was detected as described above.

### Surface plasmon resonance (SPR)

SPR assays were carried out using a BIAcore T200 instrument (GE Healthcare). Briefly, the surface of L1 sensor chip was cleaned by a 5-min injection of 40 mM octyl D-glucoside at a flow rate of 5 μl/min. Vesicles containing DOPC alone, 3% (mole/mole) of PtdIns(4,5)*P*_2_ or PtdIns(3,4,5)*P*_3_ were generated through 50 nm NanoSizer Liposome Extruders (T&T Scientific). Vesicles were immobilized on the L1 sensor chip surface, resulting in a signal of around 6500 to 8500 resonance units (RUs). Purified test proteins were injected over the surface at five or more sequentially diluted concentrations, at a flow rate of 3 μl/min. All experiments were performed at 25^◦^C in HBS buffer (pH 7.4). The sensorgrams were recorded during the association and disassociation and were analyzed using BIAevaluation software (GE Healthcare). SPR signals were corrected for background (DOPC) binding, and a binding isotherm was generated from equilibrium response (R_eq_) versus the concentration (C) of proteins. The equilibrium dissociation constant (*K*_D_) was derived from steady-state affinity analysis by nonlinear least-squares fitting of the binding isotherm using the equation R_eq_=R_max_/(1+*K*_D_/C). The percentage of maximal binding was determined at each protein concentration as equilibrium response divided by the maximum response measured at saturation.

### F-actin depolymerization assay

Actin Polymerization Biochem Kit (Cytoskeleton) was used to investigate F-actin depolymerization by cofilin in the absence or presence of control protein BSA, TNFAIP8, and SUVs containing 10% PtdIns(4,5)*P*_2_, 10% PtdIns(3,4,5)*P*_3_ or 10% PtdIns(4,5)*P*_2_ plus 10% PtdIns(3,4,5)*P*_3_, according to the manufacturer’s protocol with minor modifications. 2 mM SUVs (1 mM total available lipids) were pretreated with 11 μM control protein BSA or TNFAIP8 protein for 1 h; then 11 μM cofilin was added to each reaction and incubated for 20 min. In addition, 11 μM cofilin was also incubated with 11 μM TNFAIP8 or BSA for 20 min (control reactions). Pyrene-labeled F-actin stock was diluted to the concentration of 0.1 mg/ml with general actin buffer containing 25 mM Tris-HCl (pH 8.0) and 0.2 mM CaCl2. For each sample, 100 μl of diluted pyrene-labeled F-actin stock was used. The fluorescence signals were measured immediately before and after adding 10 μl of one of the following reagents: (a) BSA (22 μM), (b) cofilin (11 μM) + BSA (11 μM), (c) cofilin (11 μM) + TNFAIP8 (11 μM), (d) cofilin (11 μM) + BSA (11 μM) + SUVs, (e) cofilin (11 μM) + TNFAIP8 (11 μM) + SUVs, or (f) vehicle (i.e., 25 mM HEPES, 150 mM NaCl, pH 7.5). The experiments were performed in duplicates or triplicates. The fluorescence signals were detected by the Infinite 200 Pro fluorescence plate reader (Tecan). The fluorescence measurements of each sample before adding the above reagents were set as 100%. The curves were fitted to one-phase exponential decay equations (GraphPad Prism) to assess the degree of F-actin depolymerization, which was calculated as differences in fluorescence before and after adding the above reagents for each sample. Additionally, results were also presented as the differences in the remaining F-actin over a period of time. TNFAIP8 alone did not affect F-actin depolymerization, which was not shown.

### Statistics

All statistical analysis was done using GraphPad Prism 8 software. *P* values were calculated based on two-tailed, unpaired Student’s *t*-tests unless otherwise specified. *P* < 0.05 was considered statistically significant.

