## Supplementary Information for "TNFAIP8 is a novel phosphoinositide-binding inhibitory regulator of Rho GTPases that promotes cancer cell migration"

**This PDF file includes the following:**

**Supplementary Figures 1 to 9**

**Supplementary Figure Legends**

**Supplementary Tables 1 to 4**

**Supplementary Video Legends**

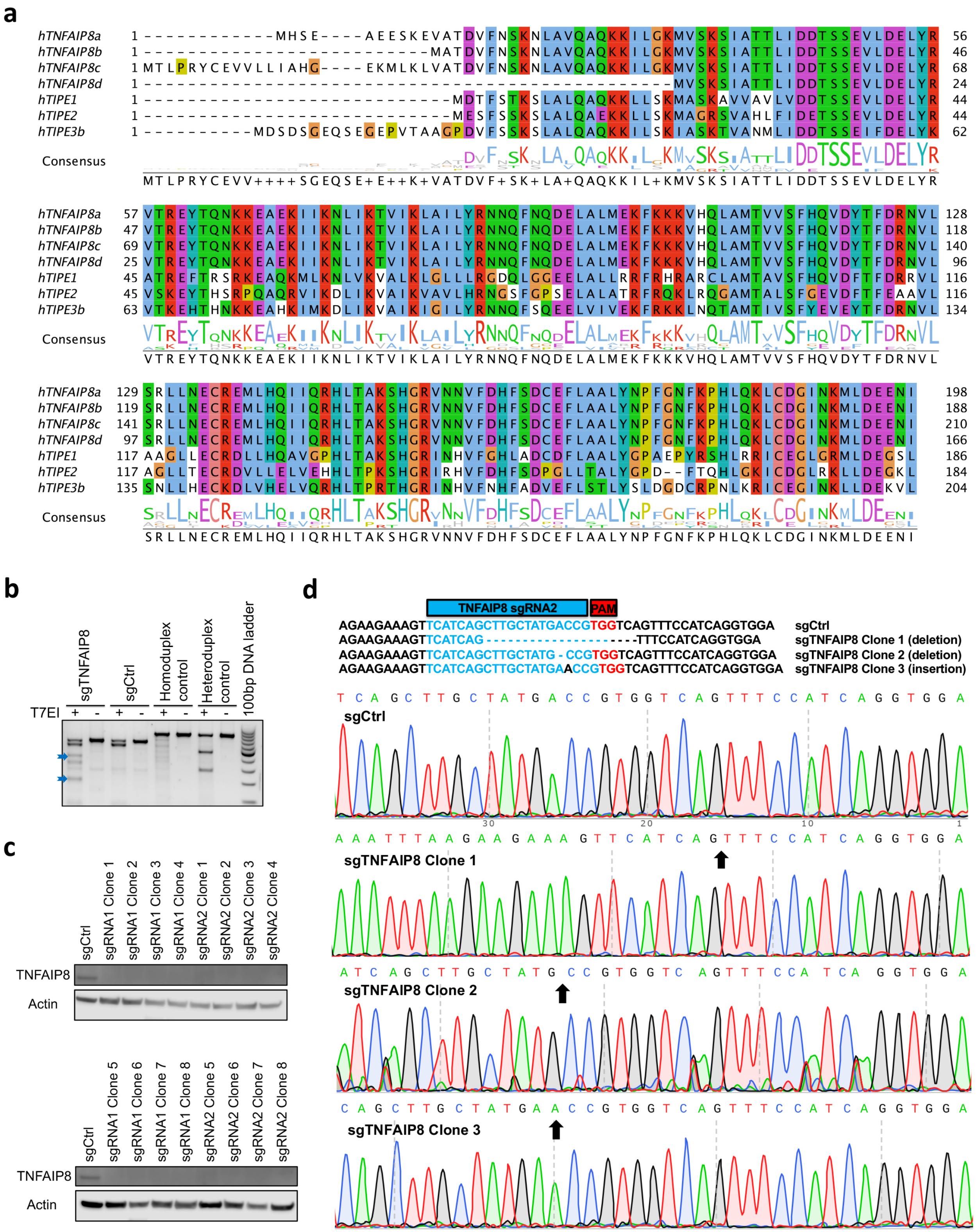
a
hTNFAIP8a 1 ------------MHSE----AEESKEVATDVFNSKNLAVQAQKKILGKMVSKSIATTLIDDTSSEVLDELYR 56
hTNFAIP8b 1 ---------------------------MATDVFNSKNLAVQAQKKILGKMVSKSIATTLIDDTSSEVLDELYR 46
hTNFAIP8c 1 MTLPRYCEVVLLIAHG----EKMLKLVATDVFNSKNLAVQAQKKILGKMVSKSIATTLIDDTSSEVLDELYR 68
hTNFAIP8d 1 ------------------------------------------------MVSKSIATTLIDDTSSEVLDELYR 24
hTIPE1 1 ----------------------------MDTFSTKSLALQAQKKLLSKMASKAVVAVLVDDTSSEVLDELYR 44
hTIPE2 1 ----------------------------MESFSSKSLALQAEKKLLSKMAGRSVAHLFIDETSSEVLDELYR 44
hTIPE3b 1 ----------MDSDSGEQSEGEPVTAAGPDVFSSKSLALQAQKKILSKIASKTVANMLIDDTSSEIFDELYR 62
Consensus
MTLPRYCEVV++++SGEQSE+E++K+VATDVF+SK+LA+QAQKKIL+KMVSKSIATTLIDDTSSEVLDELYR
hTNFAIP8a 57 VTREYTQNKKEAEKIIKNLIKTVIKLAILYRNNQFNQDELALMEKFKKKVHQLAMTVVSFHQVDYTFDRNVL 128
hTNFAIP8b 47 VTREYTQNKKEAEKIIKNLIKTVIKLAILYRNNQFNQDELALMEKFKKKVHQLAMTVVSFHQVDYTFDRNVL 118
hTNFAIP8c 69 VTREYTQNKKEAEKIIKNLIKTVIKLAILYRNNQFNQDELALMEKFKKKVHQLAMTVVSFHQVDYTFDRNVL 140
hTNFAIP8d 25 VTREYTQNKKEAEKIIKNLIKTVIKLAILYRNNQFNQDELALMEKFKKKVHQLAMTVVSFHQVDYTFDRNVL 96
hTIPE1 45 ATREFTRSRKEAQKMLKNLVKVALKLGLLLRGDQLGGEELALLRRFRHRARCLAMTAVSFHQVDFTFDRRVL 116
hTIPE2 45 VSKEYTHSRPQAQRVIKDLIKVAIKVAVLHRNGSFGPSELALATRFRQKLRQGAMTALSFGEVDFTFEAAVL 116
hTIPE3b 63 VTKEHTHNKKEAHKIMKDLIKVAIKIGILYRNNQFSQEELVIVEKFRKKLNQTAMTIVSFYEVEYTFDRNVL 134
Consensus
VTREYTQNKKEAEKIIKNLIKTVIKLAILYRNNQFNQDELALMEKFKKKVHQLAMTVVSFHQVDYTFDRNVL
hTNFAIP8a 129 SRLLNECREMLHQIIQRHLTAKSHGRVNNVFDHFSDCEFLAALYNPFGNFKPHLQKLCDGINKMLDEENI 198
hTNFAIP8b 119 SRLLNECREMLHQIIQRHLTAKSHGRVNNVFDHFSDCEFLAALYNPFGNFKPHLQKLCDGINKMLDEENI 188
hTNFAIP8c 141 SRLLNECREMLHQIIQRHLTAKSHGRVNNVFDHFSDCEFLAALYNPFGNFKPHLQKLCDGINKMLDEENI 210
hTNFAIP8d 97 SRLLNECREMLHQIIQRHLTAKSHGRVNNVFDHFSDCEFLAALYNPFGNFKPHLQKLCDGINKMLDEENI 166
hTIPE1 117 AAGLLECRDLLHQAVGPHLTAKSHGRINHVFGHLADCDFLAALYGPAEPYRSHLRRICEGLGRMLDEGSL 186
hTIPE2 117 AGLLTECRDVLLELVEHHLTPKSHGRIRHVFDHFSDPGLLTALYGPD--FTQHLGKICDGLRKLLDEGKL 184
hTIPE3b 135 SNLLHECKDLVHELVQRHLTPRTHGRINHVFNHFADVEFLSTLYSLDGDCRPNLKRICEGINKLLDEKVL 204
Consensus
SRLLNECREMLHQIIQRHLTAKSHGRVNNVFDHFSDCEFLAALYNPFGNFKPHLQKLCDGINKMLDEENI
b
sgTNFAIP8
sgCtrl
Homoduplex control
Heteroduplex control
100bp DNA ladder
T7EI + - + - + - + -
c
sgCtrl
sgRNA1 Clone 1
sgRNA1 Clone 2
sgRNA1 Clone 3
sgRNA1 Clone 4
sgRNA2 Clone 1
sgRNA2 Clone 2
sgRNA2 Clone 3
sgRNA2 Clone 4
TNFAIP8
Actin
sgCtrl
sgRNA1 Clone 5
sgRNA1 Clone 6
sgRNA1 Clone 7
sgRNA1 Clone 8
sgRNA2 Clone 5
sgRNA2 Clone 6
sgRNA2 Clone 7
sgRNA2 Clone 8
TNFAIP8
Actin
d
TNFAIP8 sgRNA2
PAM
AGAAGAAAGTTCATCAGCTTGCTATGACCGTGGTCAGTTTCCATCAGGTGGA sgCtrl
AGAAGAAAGTTCATCAG-------------------TTTCCATCAGGTGGA sgTNFAIP8 Clone 1 (deletion)
AGAAGAAAGTTCATCAGCTTGCTATG-CCGTGGTCAGTTTCCATCAGGTGGA sgTNFAIP8 Clone 2 (deletion)
AGAAGAAAGTTCATCAGCTTGCTATGAACCGTGGTCAGTTTCCATCAGGTGGA sgTNFAIP8 Clone 3 (insertion)
sgCtrl
sgTNFAIP8 Clone 1
sgTNFAIP8 Clone 2
sgTNFAIP8 Clone 3

**Supplementary Figure 1. Sequence alignment of TIPE family and validation of CRISPR/Cas9-mediated TNFAIP8 gene knockout in HL-60 cells.** (**a**) Sequence alignment of human TNFAIP8 isoforms, TIPE1, TIPE2, and TIPE3 proteins by Clustal Omega. (**b**) T7EI genomic cleavage detection (GCD) assay performed on the genomic DNA of cells harvested three days after puromycin selection. Arrows indicate digested reannealed PCR products by T7EI (which cleaved mismatched DNA heteroduplexes). (**c**,**d**) Western blot (**c**) and Sanger sequencing (**d**) validated the insertions or deletions (indels) in isolated single clone cell lines that generated truncated TNFAIP8 proteins and premature stop codons, respectively. Data are representative of two independent experiments with similar results (**b**,**c**).

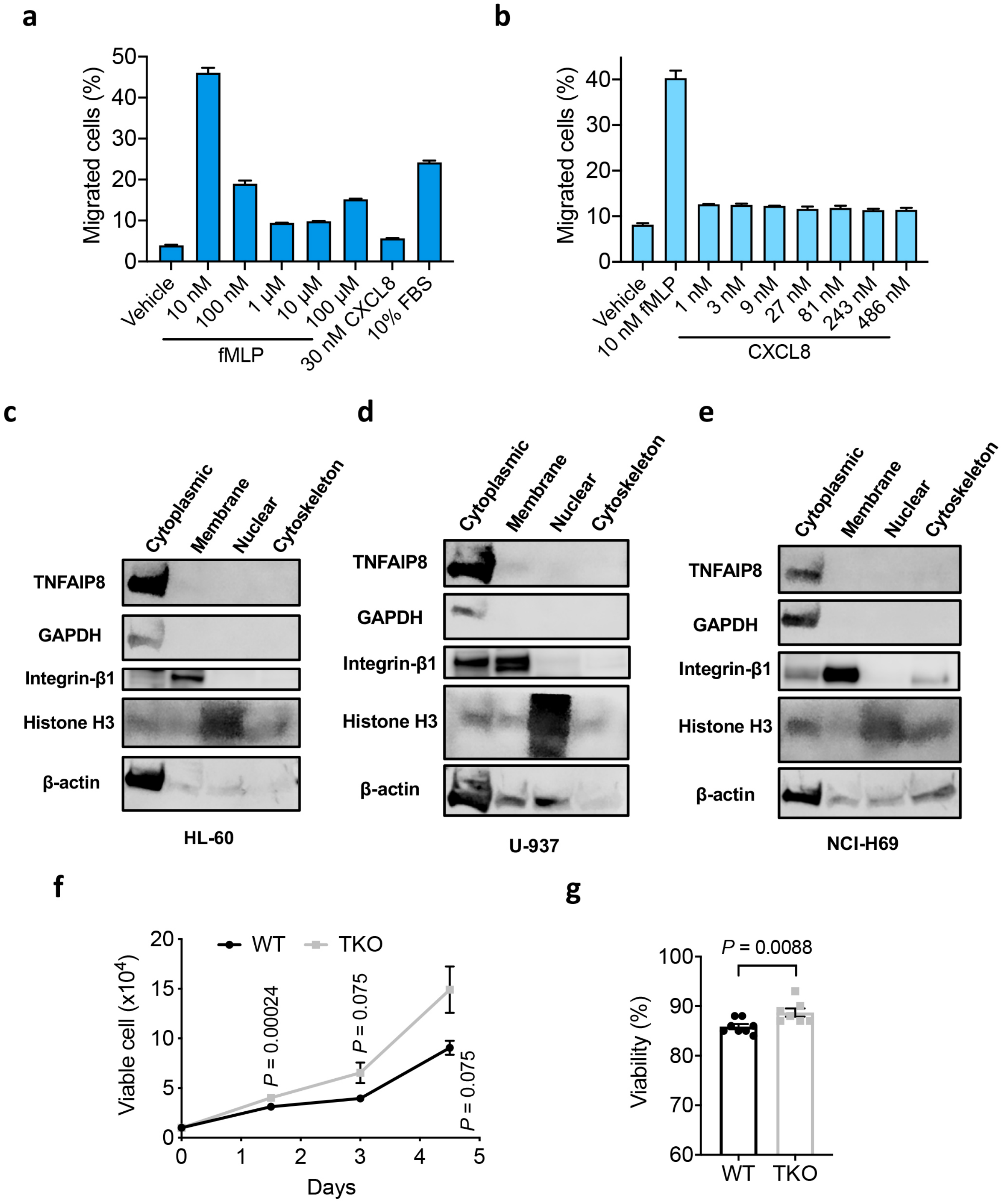

a
Migrated cells (%)
Vehicle
10 nM
100 nM
1 μM
10 μM
100 μM
30 nM CXCL8
10% FBS
fMLP
b
Migrated cells (%)
Vehicle
10 nM fMLP
1 nM
3 nM
9 nM
27 nM
81 nM
243 nM
486 nM
CXCL8
c
Cytoplasmic
Membrane
Nuclear
Cytoskeleton
TNFAIP8
GAPDH
Integrin-β1
Histone H3
β-actin
HL-60
d
U-937
e
NCI-H69
f
WT
TKO
Viable cell (x10^4)
Days
P = 0.00024
P = 0.075
P = 0.075
g
P = 0.0088
Viability (%)
WT
TKO

**Supplementary Figure 2. Stimulation of dHL-60 cells and TNFAIP8 subcellular localization in the quiescent state.** (**a**,**b**) Percentages of WT dHL-60 cells that migrated through 3 μm pore size non-coated Transwell filters towards various concentrations of fMLP (**a**) or CXCL8 (**b**). (**c**–**e**) Cytosolic localization of TNFAIP8 under resting condition in human HL-60 promyelocytic leukemia cells (**c**), U-937 lymphoma cells (**d**), and NCI-H69 small cell lung carcinoma cells (**e**). Subcellular fractionations of proteomic samples were derived by Qproteome cell compartment assay, and components from membrane, cytosol, nuclear, and cytoskeleton were analyzed by western blot using antibodies to the indicated proteins. (**f**) Viable cell numbers after inducing HL-60 differentiation for the indicated time measured by CTG assay ($n$ = 3 clones). (**g**) dHL-60 cellular viability after inducing differentiation for 5 days detected by trypan blue exclusion assay ($n$ = 8 WT or 7 TKO clones). Data are mean ± s.d. from technical duplicates (**a**,**b**) or mean ± s.e.m. (**f**,**g**), and represent two (**a**–**e**) or three (**f**,**g**) independent experiments with similar results. $P$ values are from two-sided unpaired Student's $t$-test (**f**,**g**).

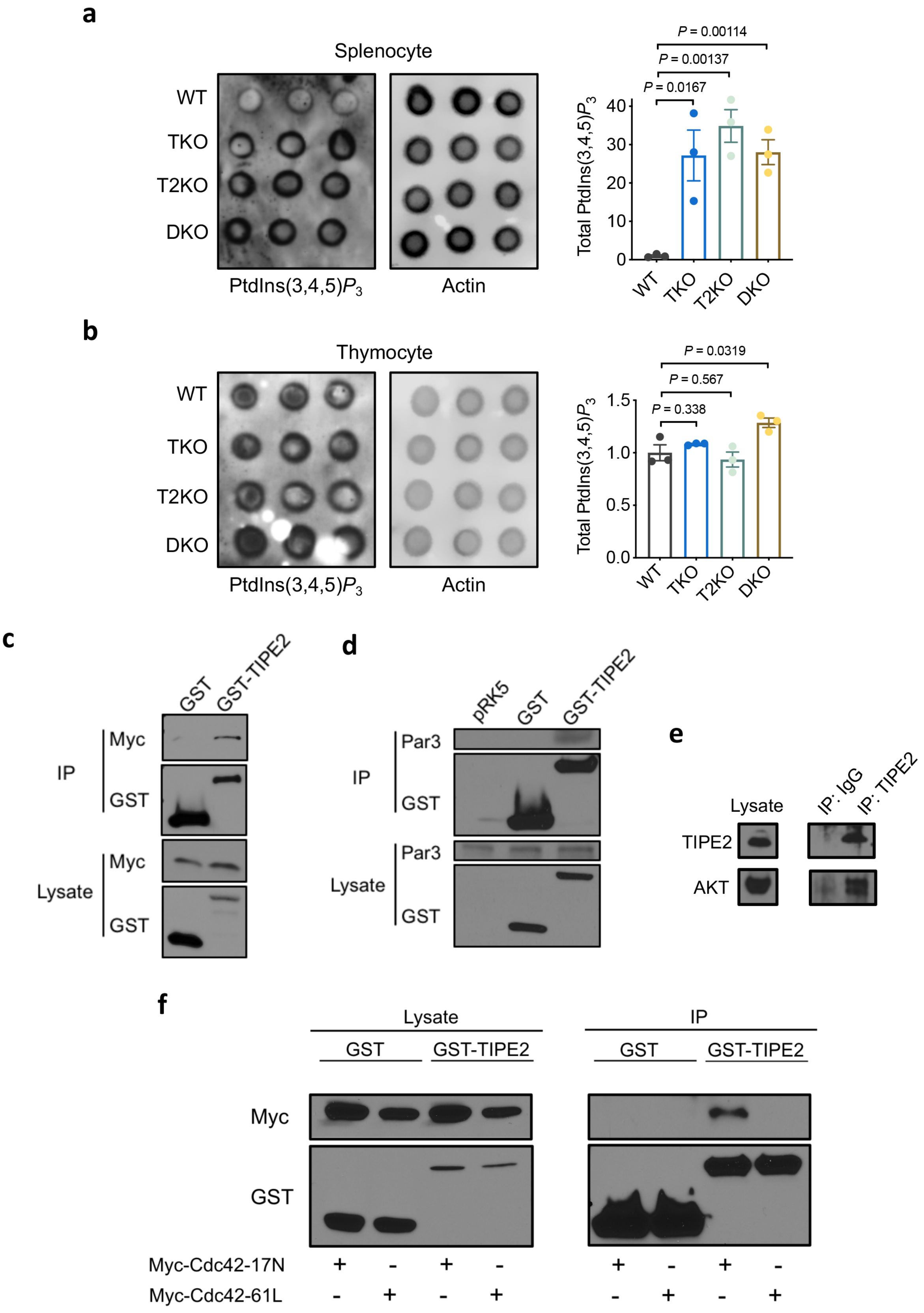
a
Splenocyte
WT
TKO
T2KO
DKO
PtdIns(3,4,5)P3
Actin
P = 0.00114
P = 0.00137
P = 0.0167
Total PtdIns(3,4,5)P3
40
30
20
10
0
WT
TKO
T2KO
DKO
b
Thymocyte
WT
TKO
T2KO
DKO
PtdIns(3,4,5)P3
Actin
P = 0.0319
P = 0.567
P = 0.338
Total PtdIns(3,4,5)P3
1.5
1.0
0.5
0.0
WT
TKO
T2KO
DKO
c
GST
GST-TIPE2
IP
Myc
GST
Lysate
Myc
GST
d
pRK5
GST
GST-TIPE2
IP
Par3
GST
Lysate
Par3
GST
e
Lysate
IP: IgG
IP: TIPE2
TIPE2
AKT
f
Lysate
IP
GST
GST-TIPE2
GST
GST-TIPE2
Myc
GST
Myc-Cdc42-17N
Myc-Cdc42-61L
+ - + -
- + - +
+ - + -
- + - +

**Supplementary Figure 3. Increased total PtdIns(3,4,5)$P_3$ level in TNFAIP8 and TIPE2-deficient murine splenocytes and thymocytes under resting condition, and co-immunoprecipitation analysis of TIPE2 interacting proteins.** (**a**,**b**) Cellular PtdIns(3,4,5)$P_3$ level in WT, *Tnfaip8*$^{-/-}$ (TKO), *Tipe2*$^{-/-}$ (T2KO), and *Tnfaip8*$^{-/-}$*Tipe2*$^{-/-}$ (DKO) murine splenocytes (**a**) or thymocytes (**b**) measured by protein-lipid overlay assay with GST-GRP1-PH protein. Signal from WT cells was set as 1 ($n$ = 3 mice per group). (**c**,**d**) Immunoblot analysis of the indicated proteins in 293T cells expressing Myc-tagged Cdc42 (**c**) or empty pRK5 vector (**d**), along with GST or recombinant GST-tagged TIPE2, assessed before (lower panels) or after (upper panels) pull-down with glutathione beads. (**e**) Lysates of Raw 264.7 cells were subjected to immunoprecipitation with anti-TIPE2 or control antibody immunoglobulin (IgG). The precipitates were analyzed by western blot using anti-TIPE2 and anti-AKT antibodies. (**f**) Immunoblot analysis of the indicated proteins in 293T cells transfected with GST or recombinant GST-tagged TIPE2, along with Myc-tagged Cdc42-17N or Cdc42-61L mutant, assessed before (left panel) or after (right panel) pull-down with glutathione beads. Data represent two (**a**,**b**,**d**,**e**) or three (**c**,**f**) independent experiments with similar results (mean ± s.e.m.). $P$ values are from two-sided unpaired Student's $t$-test (**a**,**b**).

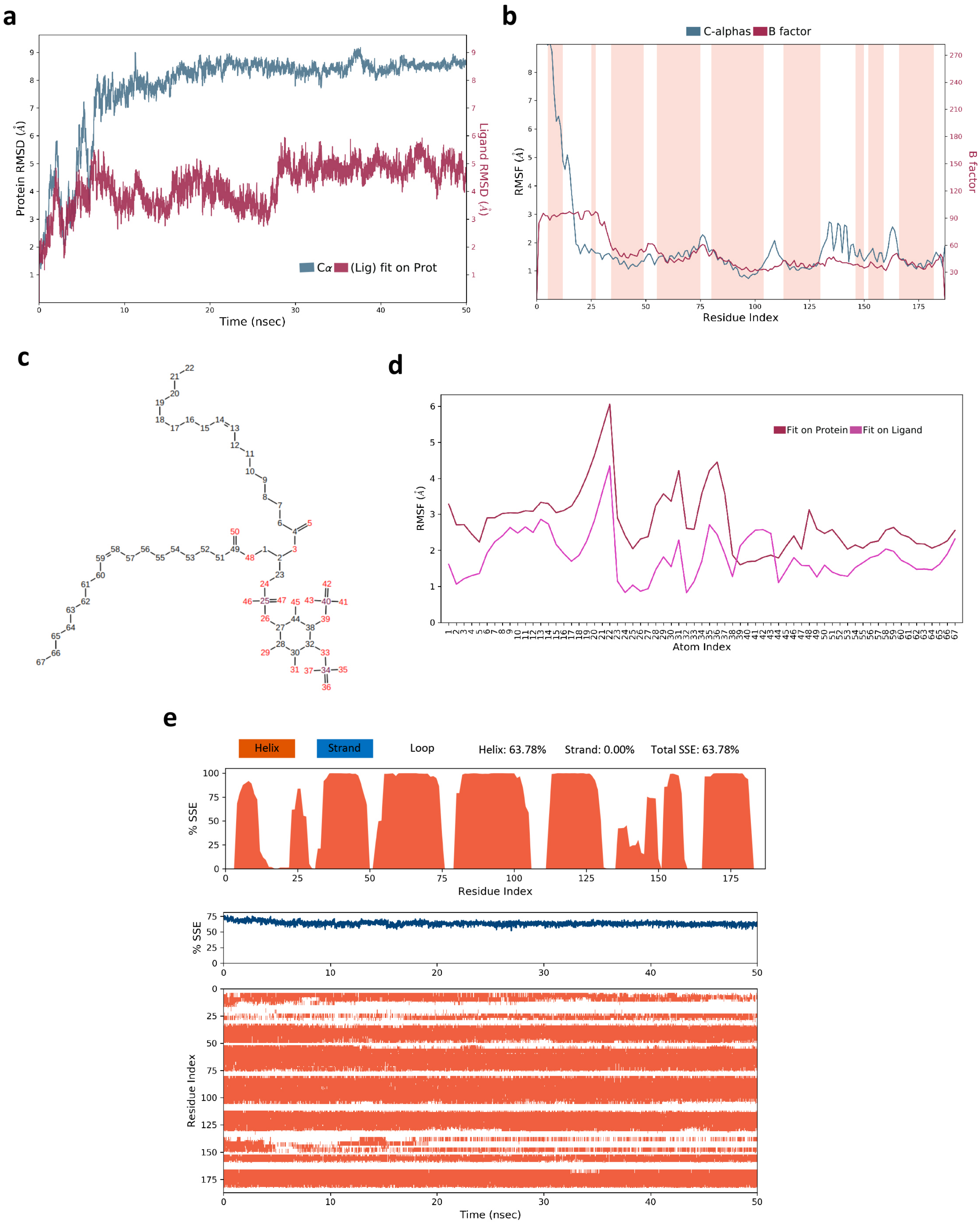

a
Protein RMSD (Å)
Ligand RMSD (Å)
Time (nsec)
Cα (Lig) fit on Prot
b
C-alphas B factor
RMSF (Å)
B factor
Residue Index
c
d
Fit on Protein Fit on Ligand
RMSF (Å)
Atom Index
e
Helix
Strand
Loop
Helix: 63.78%
Strand: 0.00%
Total SSE: 63.78%
% SSE
Residue Index
% SSE
Residue Index
Time (nsec)

**Supplementary Figure 4. Root mean square deviation (RMSD), root mean square fluctuations (RMSF), and secondary structure element (SSE) in molecular dynamics (MD) simulation of TNFAIP8 interactions with PtdIns(4,5)$P_2$.** (**a**) TNFAIP8 protein RMSD (left y-axis) and PtdIns(4,5)$P_2$ ligand RMSD (right y-axis) measuring the average changes in displacement of selected atoms with respect to the reference frame (time = 0). (**b**) RMSF for each residue in the TNFAIP8 protein chain (left y-axis) showing the time-averaged fluctuation of the square deviation of the designated residue atoms over the entire simulation time. The RMSF simulation correlated well with experimental crystallographic B factors from PDB database (right y-axis). Regions that corresponded to α helices (defined by persisting for more than 70% of the simulation) were highlighted in red. (**c**,**d**) RMSF (**d**) for each atom in the PtdIns(4,5)$P_2$ ligand (**c**) over the entire simulation time aligning on the ligand (representing internal fluctuations of PtdIns(4,5)$P_2$) or aligning on the protein (representing fluctuations with respect to the binding site). (**e**) The upper panel shows the percentage of time each residue contributed to α-helix (highlighted in red), β-strand (highlighted in blue), or loop SSE. The middle panel summarizes the SSE content for each trajectory frame over the simulation time. The bottom panel shows the SSE composition as a function of time and residue index.

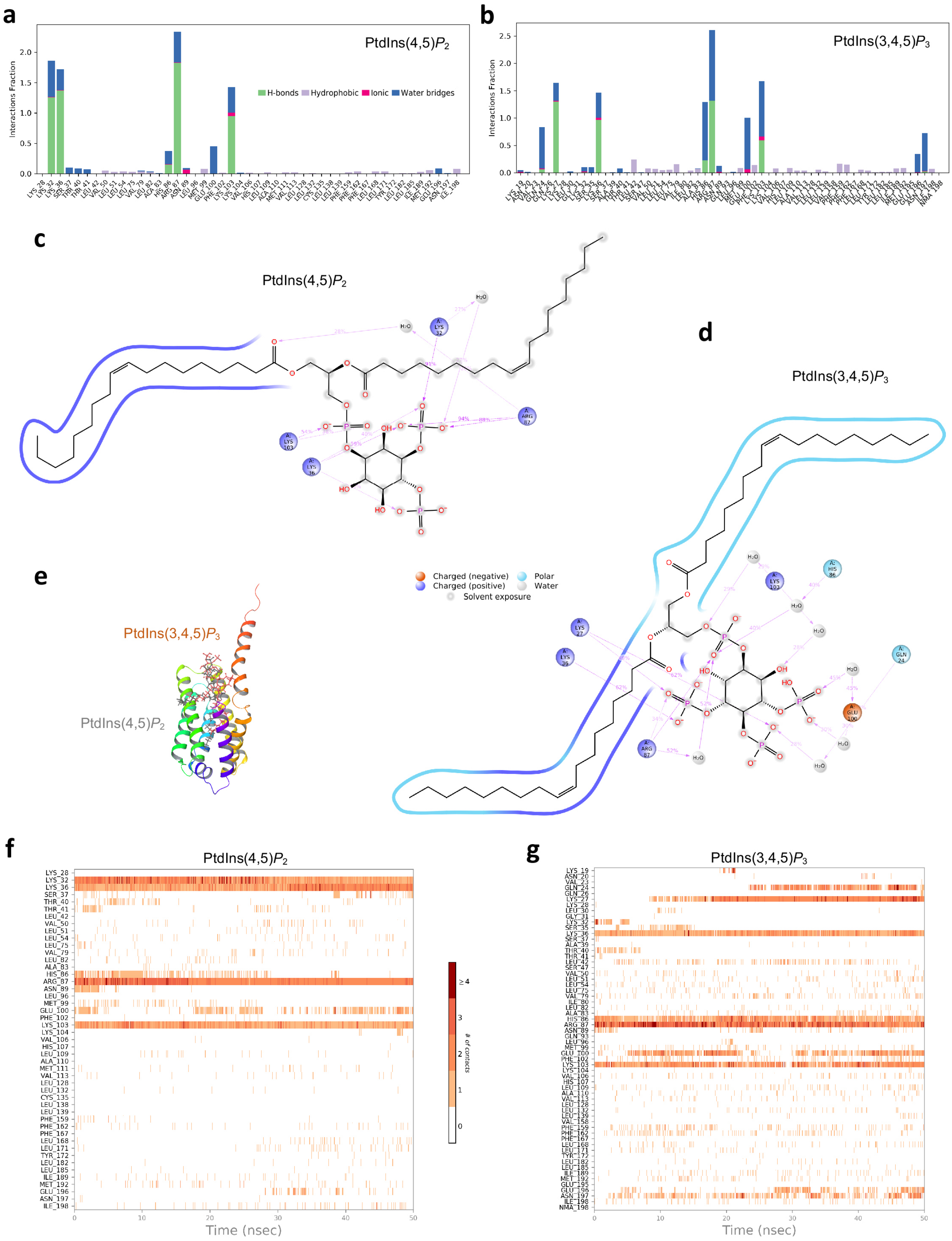
a
PtdIns(4,5)$P_2$
Interactions Fraction
H-bonds
Hydrophobic
Ionic
Water bridges
b
PtdIns(3,4,5)$P_3$
Interactions Fraction
c
PtdIns(4,5)$P_2$
d
PtdIns(3,4,5)$P_3$
e
PtdIns(3,4,5)$P_3$
PtdIns(4,5)$P_2$
Charged (negative)
Charged (positive)
Polar
Water
Solvent exposure
f
PtdIns(4,5)$P_2$
Time (nsec)
### of contacts
g
PtdIns(3,4,5)$P_3$
Time (nsec)

**Supplementary Figure 5. Protein-ligand contacts and ligand-protein contacts in MD simulations of TNFAIP8 interactions with PtdIns(4,5)$\boldsymbol{P_2}$ or PtdIns(3,4,5)$\boldsymbol{P_3}$.** (**a**,**b**) Bar charts showing the fraction of simulation time that the PtdIns(4,5)$P_2$ (**a**) or PtdIns(3,4,5)$P_3$ (**b**) ligand was in contact with various protein residues, including H-bonds, hydrophobic, ionic, and water bridges. (**c**,**d**) Ligand interaction diagrams of PtdIns(4,5)$P_2$ (**c**) and PtdIns(3,4,5)$P_3$ (**d**) atom interactions with neighboring protein residues and other species (marked with spheres) at the TNFAIP8 binding sites. Interactions that occurred more than 25% of the 50 ns simulation time in the entire trajectory are shown. (**e**) Cartoon presentation of TNFAIP8 in complex with PtdIns(4,5)$P_2$ (in grey) or PtdIns(3,4,5)$P_3$ (in orange) shown by superimposed stick models. (**f**,**g**) The number of interactions for each protein residue in contact with the bound ligand PtdIns(4,5)$P_2$ (**f**) or PtdIns(3,4,5)$P_3$ (**g**) as a function of simulation time (color-coded in shades of orange).

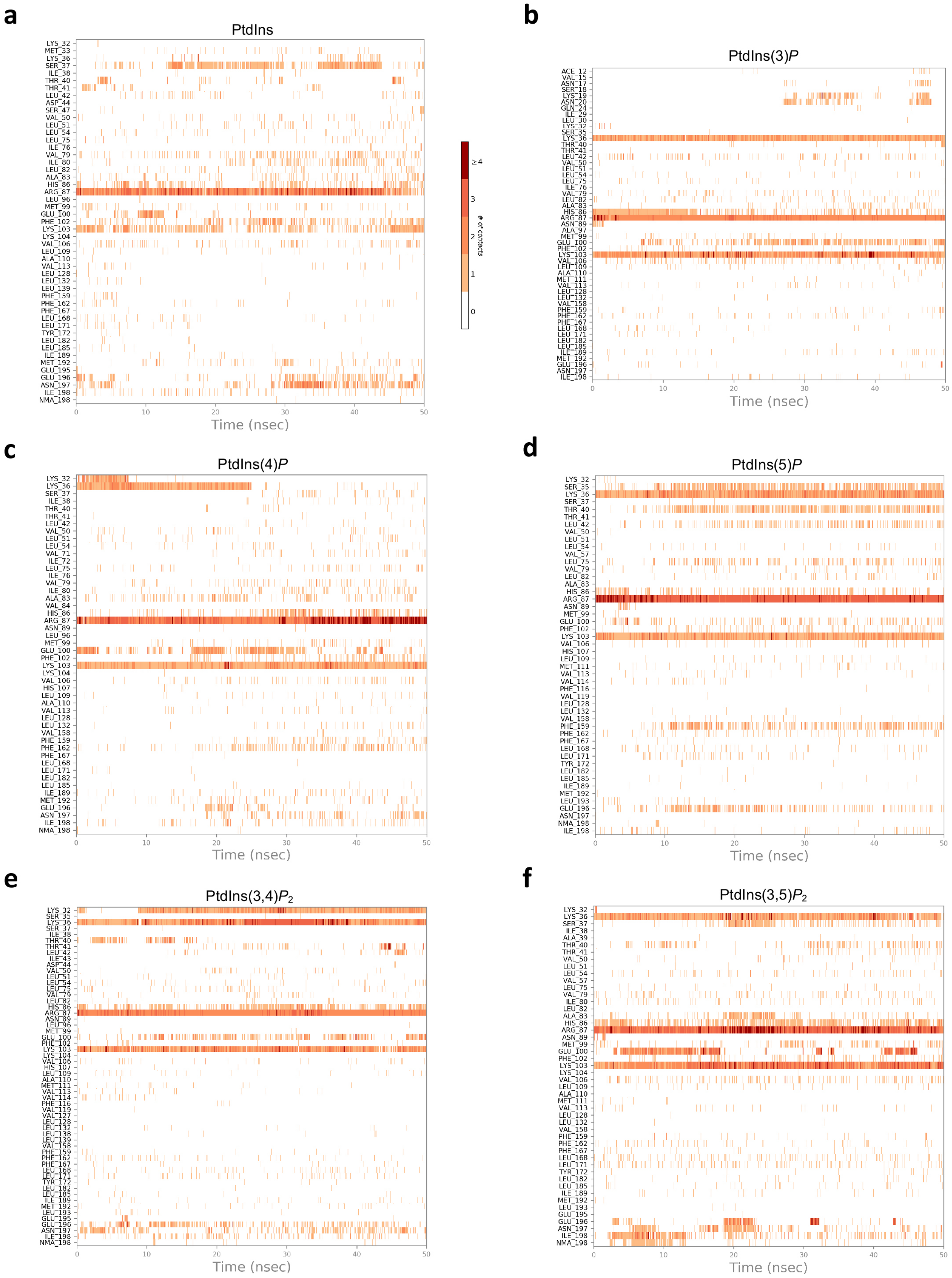

a
PtdIns
b
PtdIns(3)P
c
PtdIns(4)P
d
PtdIns(5)P
e
PtdIns(3,4)P2
f
PtdIns(3,5)P2
Time (nsec)
### of contacts
≥4
3
2
1
0

**Supplementary Figure 6. Protein-ligand contacts in MD simulations of TNFAIP8 interactions with phosphatidylinositol, monophosphates, or bisphosphates.** (**a**–**f**) The number of interactions for each protein residue in contact with PtdIns (**a**), PtdIns(3)*P* (**b**), PtdIns(4)*P* (**c**), PtdIns(5)*P* (**d**), PtdIns(3,4)$P_2$ (**e**), or PtdIns(3,5)$P_2$ (**f**) as a function of simulation time (color-coded in shades of orange).

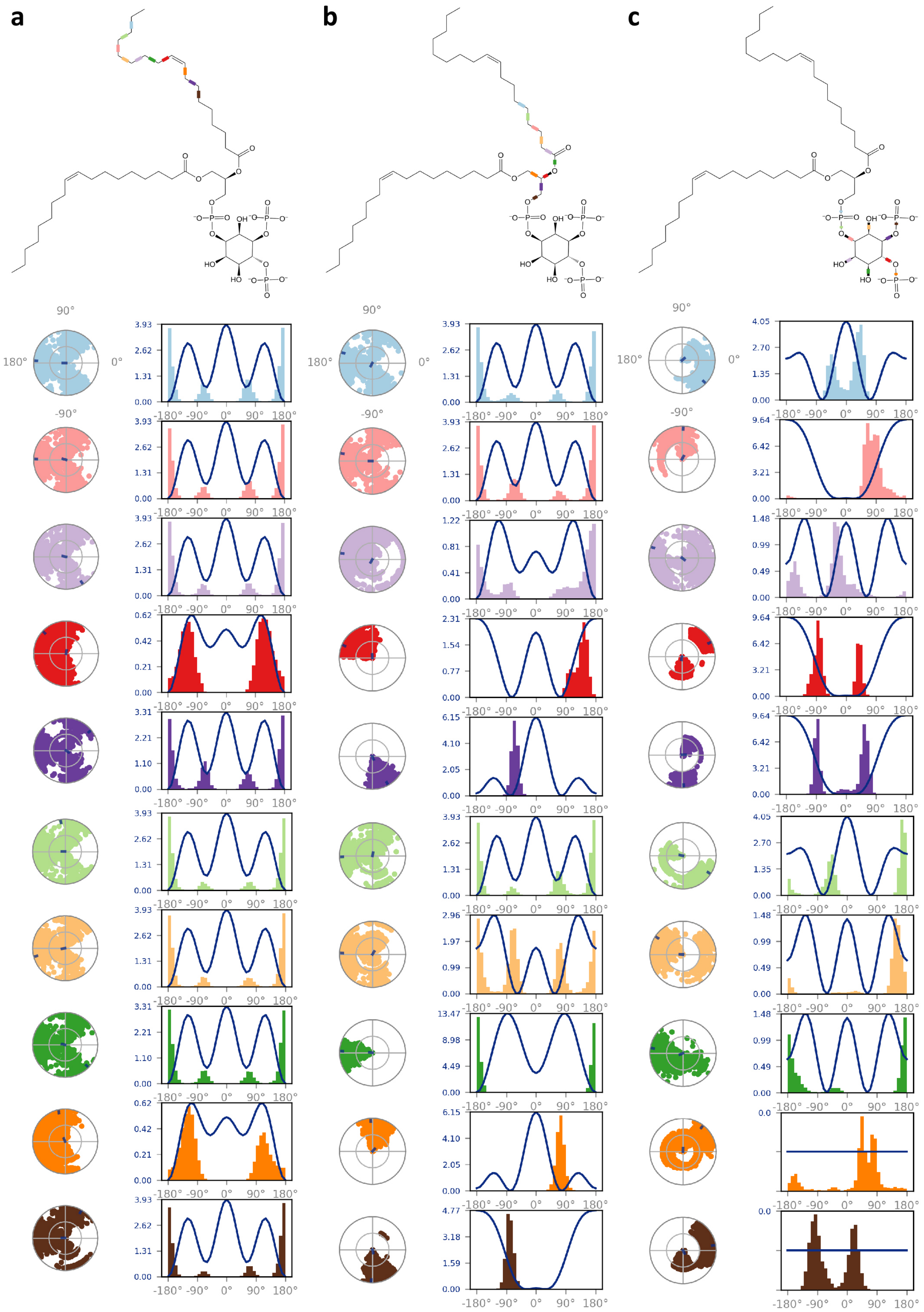

a
b
c

**Supplementary Figure 7. Ligand torsion charts summarizing the torsional conformation of rotatable bonds in PtdIns(4,5)$\boldsymbol{P_2}$ throughout the 50 ns simulation trajectory.** (**a**–**c**) The top panels show the schematic PtdIns(4,5)$P_2$ ligand with color-coded rotatable bonds, and the bottom panels show the dial plots and bar plots of the same color coding for each bond torsion on the structure. The dial plots display the conformation of the torsion as a function of time, where the beginning of the simulation was in the center and the time evolution was plotted radially outwards, and the angular coordinate was the torsional angle. The bar plots show the probability density of the torsions as a function of angle (representing the average over the simulation time). The bar plots also describe the torsional potential information of the rotatable bond in kcal/mol (left y-axis).

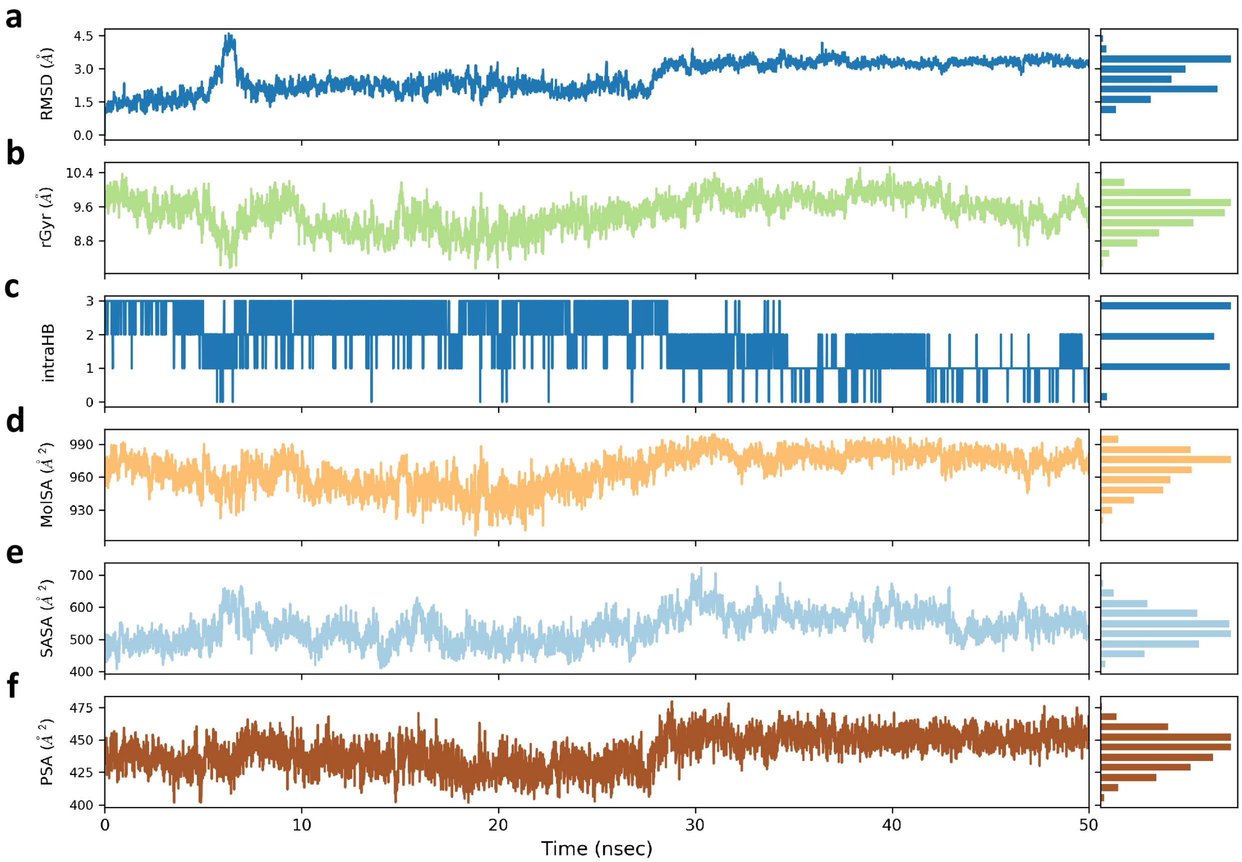


**Supplementary Figure 8. Ligand properties of PtdIns(4,5)$P_2$ throughout the 50 ns simulation trajectory.** (**a**–**f**) Ligand properties including RMSD with respect to the initial conformation (**a**), radius of gyration (rGyr) (**b**), number of intramolecular hydrogen bonds (intraHB) (**c**), molecular surface area (MolSA) (**d**), solvent-accessible surface area (SASA) (**e**), and polar surface area (PSA) (**f**). The left charts display the values of the property as a function of time. The right bar plots show the proportion of time spent in each of ten value ranges (equally divided over the range of property values).

a

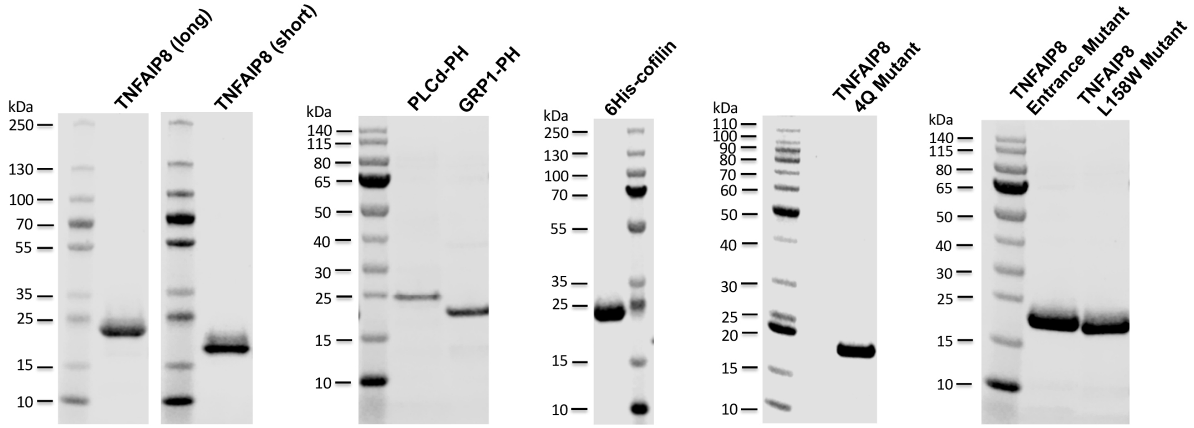


b

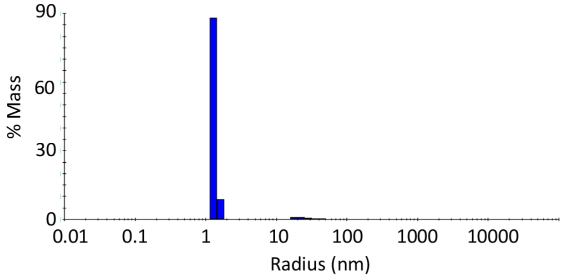


c

| Cumulant fit | | | |
|---|---|---|---|
| Radius (nm) | %PD | Baseline | SOS |
| 32.3 | 18.5 | 1 | 7.971 |
| **Regularization fit** | | | |
| Peak | Radius (nm) | %PD | %Mass |
| Peak 1 | 1.3 | 7.5 | 96.8 |
| Peak 2 | 27.1 | 41.52 | 3.2 |

d e f

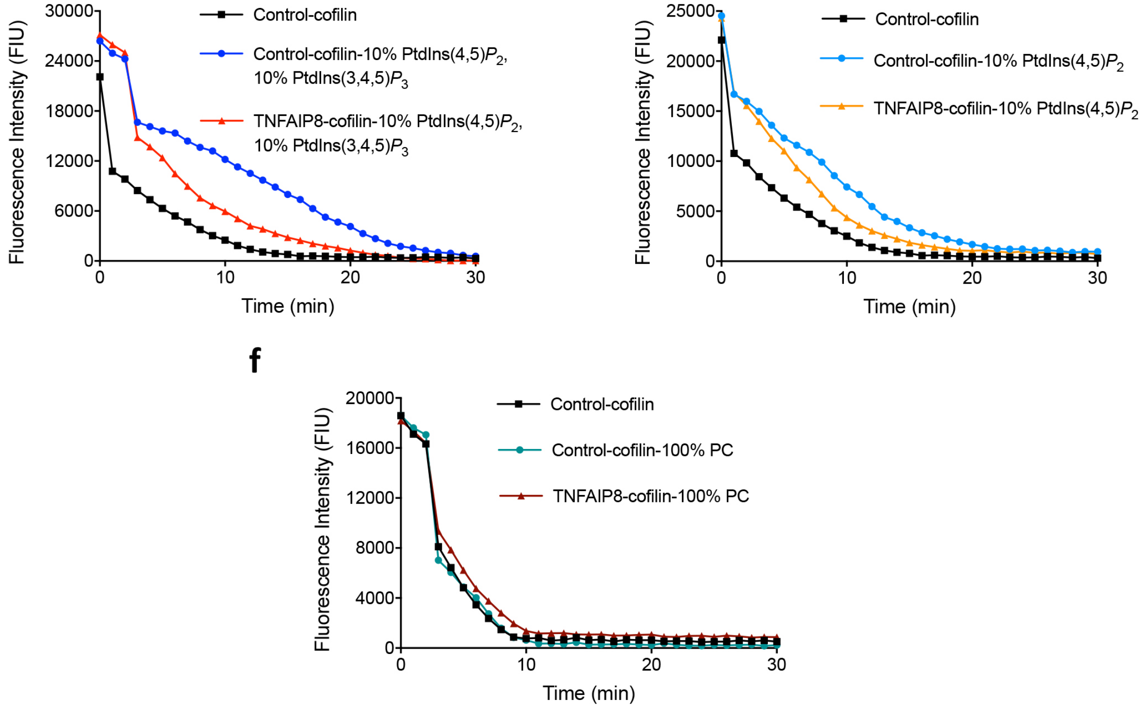

**Supplementary Figure 9. Protein purification with pET-SUMO system and effects of TNFAIP8 on cofilin-dependent F-actin depolymerization *in vitro*.** (**a**) Recombinant 6His-SUMO- or 6His-tagged proteins were expressed from *Escherichia coli* BL21(DE3) and purified using Ni-NTA resin. Final eluates were separated by SDS–PAGE and stained with Coomassie Blue G-250. Purified proteins were at least 95% pure judging from overloaded Coomassie Blue stained SDS gels. (**b**,**c**) The size distribution of hydrodynamics radius of purified TNFAIP8 in solution measured by DynaPro NanoStar (**b**), and the data fitting of dynamic light scattering (DLS) (**c**). (**d**–**f**) Representative time course of cofilin-dependent F-actin depolymerization performed in the presence or absence of control protein (**d**–**f**), TNFAIP8 (**d**–**f**), SUVs containing 10% PtdIns(4,5)$P_2$/10% PtdIns(3,4,5)$P_3$ (**d**), 10% PtdIns(4,5)$P_2$ (**e**), or 100% PC (**f**). Data is shown as the difference in pyrene labeled F-actin fluorescence intensity units (FIU) over the indicated time. Data represent three independent experiments with similar results (**a**–**f**).

#### Supplementary Table 1. CRISPR sequences targeting the shared exon of all human TNFAIP8 isoforms

| Number | Sequence | PAM | Direction |
|---|---|---|---|
| 1 | 5'- GGATAACACATTCCGGTCAA -3' | AGG | (-) |
| 2 | 5'- TCATCAGCTTGCTATGACCG -3' | TGG | (+) |
| 3 | 5'- AGGTGGATTATACCTTTGAC -3' | CGG | (+) |

#### Supplementary Table 2. Primers for cloning the targeting sequences into LentiCRISPRv2 backbone and for genomic cleavage detection (GCD) CRISPR validation assay

| Primer | Forward | Reverse | GCD band size (bp) |
|---|---|---|---|
| LentiCRISPRv2 TNFAIP8 sgRNA1 | CACCGGATAACACATTCCGGTCAA | AAACTTGACCGGAATGTGTTATCC | NA |
| LentiCRISPRv2 TNFAIP8 sgRNA2 | CACCGTCATCAGCTTGCTATGACCG | AAACCGGTCATAGCAAGCTGATGAC | NA |
| LentiCRISPRv2 TNFAIP8 sgRNA3 | CACCGAGGTGGATTATACCTTTGAC | AAACGTCAAAGGTATAATCCACCTC | NA |
| sgRNA1 GCD Primer set1 | TTTAGTTCGCTTCACTTG | ACCATCACATAGTTTTTG | 600 [406, 194] |
| sgRNA1 GCD Primer set2 | TTAATTCTTTTCCTCTCTTTTT | GGTTTAAAATTCCCAAAA | 540 [371, 169] |
| sgRNA2 GCD Primer set1 | CAAATGGTGGGAAAATGAGG | GGATTATACAAGGCAGCCAA | 603 [416, 187] |
| sgRNA2 GCD Primer set2 | TTAGTTCGCTTCACTTGCTG | GGATTATACAAGGCAGCCAA | 556 [369, 187] |
| sgRNA3 GCD Primer set1 | ATACCTGTTTTTAGTTCGCTTC | TAGTTTTTGTAAGTGGGG | 600 [414, 186] |
| sgRNA3 GCD Primer set2 | TTAATTCTTTTCCTCTCTTTTT | GGTTTAAAATTCCCAAAA | 540 [371, 169] |

#### Supplementary Table 3. Glide extra precision (XP) docking scores of TNFAIP8 with various phospholipids

| Title | DOPE | PtdIns | PtdIns(3)*P* | PtdIns(4)*P* | PtdIns(5)*P* | PtdIns(3,4)$P_2$ | PtdIns(3,5)$P_2$ | PtdIns(4,5)$P_2$ | PtdIns(3,4,5)$P_3$ |
|---|---|---|---|---|---|---|---|---|---|
| XP GScore | -8.565 | -12.538 | -12.103 | -10.653 | -11.425 | -10.566 | -10.381 | -11.159 | -9.983 |
| XP LipophilicEvdW | -6.335 | -7.226 | -6.985 | -5.263 | -6.021 | -6.005 | -7.020 | -5.635 | -4.858 |
| XP PhobEn | -2.562 | -2.225 | -2.372 | -1.932 | -2.042 | -2.273 | -1.726 | -1.397 | -0.881 |
| XP Electro | -0.888 | -0.624 | -1.112 | -0.812 | -1.182 | -1.396 | -1.126 | -1.005 | -1.521 |
| XP HBond | -1.354 | -2.820 | -1.799 | -2.593 | -2.830 | -3.384 | -2.205 | -3.289 | -2.624 |
| XP RotPenal | 0.554 | 0.389 | 0.324 | 0.324 | 0.324 | 0.273 | 0.273 | 0.273 | 0.240 |
| XP ExposPenal | 2.110 | 0.397 | 0.199 | 1.123 | 0.336 | 2.367 | 1.545 | 1.393 | 1.161 |
| glide ligand efficiency | -0.168 | -0.213 | -0.190 | -0.167 | -0.180 | -0.155 | -0.152 | -0.163 | -0.136 |
| glide ligand efficiency sa | -0.622 | -0.827 | -0.758 | -0.666 | -0.715 | -0.628 | -0.616 | -0.664 | -0.564 |
| glide ligand efficiency ln | -1.736 | -2.469 | -2.333 | -2.051 | -2.201 | -1.990 | -1.953 | -2.103 | -1.837 |
| glide evdw | -53.537 | -53.920 | -51.748 | -48.131 | -56.252 | -57.685 | -56.196 | -49.237 | -43.483 |
| glide ecoul | -11.835 | -8.320 | -14.821 | -10.822 | -15.758 | -18.620 | -15.018 | -13.395 | -20.281 |
| glide energy | -65.372 | -62.240 | -66.568 | -58.953 | -72.011 | -76.304 | -71.215 | -62.632 | -63.764 |
| glide einternal | 19.848 | 23.237 | 14.926 | 0 | 25.414 | 32.837 | 35.458 | 25.322 | 0 |
| glide emodel | -82.296 | -80.284 | -93.126 | -71.798 | -87.253 | -100.966 | -78.964 | -69.301 | -91.397 |
| glide posenum | 2 | 16 | 12 | 5 | 1 | 1 | 1 | 1 | 11 |
| glide eff state penalty | 0.005 | 0.0002 | 0.104 | 0.106 | 0.107 | 0.209 | 0.214 | 0.212 | 0.317 |
| res:A103 hbond | 0 | 0 | 0 | 0 | 0 | -0.418 | -0.509 | 0 | 0 |
| res:A87 hbond | 0 | -0.500 | 0 | -0.433 | 0 | -0.083 | -0.320 | -0.159 | -0.206 |
| res:A86 hbond | 0 | 0 | 0 | 0 | 0 | -0.024 | -0.500 | -0.500 | 0 |
| res:A36 hbond | 0 | 0 | 0 | 0 | -0.555 | 0 | 0 | 0 | 0 |

#### Supplementary Table 4. Ligand binding and strain energies of various phospholipids with TNFAIP8 calculated using Prime MM-GBSA method

| Title | DOPE | PtdIns | PtdIns(3)*P* | PtdIns(4)*P* | PtdIns(5)*P* | PtdIns(3,4)$P_2$ | PtdIns(3,5)$P_2$ | PtdIns(4,5)$P_2$ | PtdIns(3,4,5)$P_3$ |
|---|---|---|---|---|---|---|---|---|---|
| Prime Coulomb | -6029 | -6012 | -6033 | -6024 | -6053 | -5959 | -5980 | -5985 | -6001 |
| Prime Covalent | 879.0 | 890.3 | 877.2 | 891.2 | 879.7 | 886.8 | 887.1 | 880.7 | 881.7 |
| Prime vdW | -888.0 | -903.2 | -890.7 | -900.0 | -902.8 | -900.9 | -901.1 | -901.1 | -898.1 |
| Prime Lipo | -412.3 | -423.5 | -412.7 | -410.2 | -412.4 | -415.6 | -411.2 | -411.1 | -403.9 |
| Prime Solv GB | -1637 | -1611 | -1663 | -1661 | -1653 | -1811 | -1810 | -1794 | -1834 |
| Prime Energy | -8247 | -8219 | -8283 | -8266 | -8303 | -8363 | -8379 | -8373 | -8420 |
| Prime Hbond | -116.1 | -115.5 | -116.9 | -116.9 | -118.1 | -118.2 | -118.9 | -118.6 | -120.2 |
| Prime MMGBSA ligand efficiency | -1.270 | -1.463 | -0.883 | -0.983 | -1.227 | -1.213 | -0.885 | -0.904 | -0.684 |
| Prime MMGBSA ligand efficiency sa | -1.905 | -2.194 | -1.325 | -1.474 | -1.840 | -1.820 | -1.327 | -1.357 | -1.026 |
| Prime MMGBSA ligand efficiency ln | -13.13 | -17.00 | -10.82 | -12.04 | -15.03 | -15.62 | -11.39 | -11.64 | -9.23 |
| MMGBSA dG Bind | -64.77 | -86.31 | -55.65 | -61.93 | -77.29 | -81.27 | -59.26 | -60.59 | -48.57 |
| MMGBSA dG Bind Coulomb | -16.24 | -51.41 | -78.31 | -87.32 | -99.78 | -111.80 | -98.66 | -122.45 | -177.84 |
| MMGBSA dG Bind Covalent | 1.73 | 11.37 | 7.93 | 9.56 | 1.48 | 15.30 | 9.90 | 10.10 | -1.89 |
| MMGBSA dG Bind Hbond | -3.531 | -2.867 | -4.311 | -4.305 | -5.443 | -5.635 | -6.318 | -5.964 | -7.551 |
| MMGBSA dG Bind Lipo | -22.65 | -33.56 | -25.52 | -22.26 | -23.49 | -35.37 | -22.02 | -18.79 | -10.69 |
| MMGBSA dG Bind Solv GB | 36.39 | 68.82 | 109.90 | 114.49 | 123.01 | 136.24 | 127.39 | 148.26 | 205.40 |
| MMGBSA dG Bind vdW | -60.47 | -78.66 | -65.33 | -72.09 | -73.07 | -80.01 | -69.55 | -71.75 | -55.99 |
| MMGBSA dG Bind(NS) | -88.72 | -114.94 | -86.46 | -82.97 | -97.47 | -105.17 | -89.51 | -96.50 | -70.03 |
| MMGBSA dG Bind(NS) Coulomb | -15.63 | -56.66 | -83.55 | -85.31 | -101.94 | -111.70 | -106.31 | -125.24 | -188.74 |
| MMGBSA dG Bind(NS) Covalent | 0 | 4.55E-13 | 0 | 4.55E-13 | -1.14E-13 | -1.14E-13 | 1.14E-13 | 1.14E-13 | -1.14E-13 |

| Title | DOPE | PtdIns | PtdIns(3)*P* | PtdIns(4)*P* | PtdIns(5)*P* | PtdIns(3,4)$P_2$ | PtdIns(3,5)$P_2$ | PtdIns(4,5)$P_2$ | PtdIns(3,4,5)$P_3$ |
|---|---|---|---|---|---|---|---|---|---|
| MMGBSA dG Bind(NS) Hbond | -3.531 | -2.867 | -4.311 | -4.305 | -5.443 | -5.635 | -6.318 | -5.964 | -7.551 |
| MMGBSA dG Bind(NS) Lipo | -34.01 | -39.62 | -34.36 | -29.16 | -33.10 | -37.03 | -29.59 | -31.35 | -22.40 |
| MMGBSA dG Bind(NS) Solv GB | 39.46 | 73.09 | 118.97 | 117.48 | 132.52 | 138.12 | 140.13 | 149.02 | 220.54 |
| MMGBSA dG Bind(NS) vdW | -75.00 | -88.88 | -83.20 | -81.68 | -89.51 | -88.92 | -87.42 | -82.97 | -71.89 |
| Lig Strain Energy | 23.95 | 28.63 | 30.80 | 21.04 | 20.18 | 23.90 | 30.24 | 35.91 | 21.46 |
| Lig Strain Coulomb | -0.614 | 5.249 | 5.243 | -2.013 | 2.156 | -0.094 | 7.654 | 2.792 | 10.893 |
| Lig Strain Covalent | 1.733 | 11.375 | 7.929 | 9.556 | 1.480 | 15.296 | 9.899 | 10.101 | -1.887 |
| Lig Strain Lipo | 11.360 | 6.059 | 8.837 | 6.903 | 9.614 | 1.657 | 7.567 | 12.565 | 11.706 |
| Lig Strain Solv GB | -3.067 | -4.275 | -9.072 | -2.993 | -9.508 | -1.879 | -12.746 | -0.768 | -15.149 |
| Lig Strain vdW | 14.535 | 10.225 | 17.867 | 9.589 | 16.438 | 8.916 | 17.871 | 11.223 | 15.898 |
| Ligand Energy | -81.327 | -31.526 | -126.436 | -102.616 | -124.905 | -180.366 | -218.230 | -211.239 | -270.192 |
| Ligand Coulomb | -44.926 | 6.997 | 13.111 | 30.898 | 14.938 | 120.170 | 86.167 | 105.608 | 144.891 |
| Ligand Covalent | 14.713 | 16.386 | 6.735 | 19.116 | 15.708 | 8.965 | 14.635 | 8.006 | 21.012 |
| Ligand Lipo | -17.749 | -18.040 | -15.240 | -15.966 | -16.975 | -8.236 | -17.222 | -20.357 | -21.252 |
| Ligand Solv GB | -24.55 | -31.05 | -124.34 | -127.40 | -127.56 | -299.09 | -288.98 | -293.79 | -391.43 |
| Ligand vdW | -8.812 | -5.822 | -6.699 | -9.266 | -11.015 | -2.171 | -12.832 | -10.708 | -23.414 |
| Complex Energy | -8247 | -8219 | -8283 | -8266 | -8303 | -8363 | -8379 | -8373 | -8420 |
| Complex Coulomb | -6029 | -6012 | -6033 | -6024 | -6053 | -5959 | -5980 | -5985 | -6001 |
| Complex Covalent | 879.0 | 890.3 | 877.2 | 891.2 | 879.7 | 886.8 | 887.1 | 880.7 | 881.7 |
| Complex Hbond | -116.1 | -115.5 | -116.9 | -116.9 | -118.1 | -118.2 | -118.9 | -118.6 | -120.2 |
| Complex Lipo | -412.3 | -423.5 | -412.7 | -410.2 | -412.4 | -415.6 | -411.2 | -411.1 | -403.9 |
| Complex Solv GB | -1637 | -1611 | -1663 | -1661 | -1653 | -1811 | -1810 | -1794 | -1834 |
| Complex vdW | -888.0 | -903.2 | -890.7 | -900.0 | -902.8 | -900.9 | -901.1 | -901.1 | -898.1 |

**Supplementary Video 1. Overall structure of TNFAIP8 C165S (PDB accession 5JXD) in complex with a phospholipid shown by ball model.** Helices are rainbow colored from $\alpha0$ (in blue) to $\alpha6$ (in red). The surface of the centrally located hydrophobic cavity is shown in grey.

**Supplementary Video 2. Representation of mutated TNFAIP8 residues to induce steric blockade in the cavity.** The structure of TNFAIP8 is represented by a cartoon model with the side chains of wall residues comprising the pocket (in yellow) shown as stick models. Four conserved residues L44 ($\alpha1$), L65 ($\alpha2$), L99 ($\alpha3$), and L158 ($\alpha5$) (in magenta) were mutated to Ws. The cavity surface is colored grey.

**Supplementary Video 3. Representation of replaced TNFAIP8 residues to generate 2Q, 4Q, and Entrance mutants.** Two positively charged residues of $\alpha0$ helix K17 and K18 (in orange) were mutated to Qs for the 2Q mutant, and two additional residues K22 and K26 (in orange) were replaced by Qs to construct the 4Q mutant. Three positively charged amino acids H76, R77, and K93 (in blue) positioned at the opening of the cavity were mutated to Qs for the Entrance mutation. The phospholipid and mutated residues are shown in stick models and cavity surface is colored grey.
